# BioPathfinder: Evidence-guided multi-agent platform enables hypothesis discovery for CAR-T engineering

**DOI:** 10.64898/2026.07.15.738646

**Authors:** Shubing Wang, Yan-Ruide Li, Qian Wang, Youcheng Yang, Xinyuan Shen, Haoran Li, Haochen Nan, Zhoulin Chen, Yichen Zhu, Bo Zhang, Haoyan Ding, Jennifer Soto, Soomin Park, Yi Zheng, Xiangliang Huang, Debiao Li, Song Li

**Author notes:** **Correspondence and requests for materials** Debiao Li, Ph.D., Biomedical Imaging Research Institute, Cedars-Sinai Medical Center, Los Angeles, CA 90048, USA., Yan-Ruide Li, Ph.D., Department of Microbiology, Immunology & Molecular Genetics, University of California, Los Angeles, Los Angeles, CA 90095, USA., Song Li, Ph.D., Department of Bioengineering University of California, Los Angeles, Los Angeles, CA 90095, USA. These authors contributed equally: Shubing Wang, Yan-Ruide Li, Qian Wang.

## Abstract

Clinical studies of chimeric antigen receptor (CAR)-T therapy generate diverse molecular and clinical evidence that remains fragmented across publications, public repositories and patient-derived datasets, limiting systematic therapeutic discovery. Here we present BioPathfinder, an evidence-guided multi-agent workflow for closed-loop biomedical discovery. BioPathfinder constructs a provenance-aware knowledge resource linking publications with patient single-cell RNA sequencing (scRNA-seq) datasets and clinical metadata, and uses role-specialized large language model agents to generate, review and prioritize diverse, falsifiable and dataset-aware mechanistic hypotheses for computational and experimental validation. Applied to a curated corpus of CAR-T-treated patient studies and matched scRNA-seq datasets, BioPathfinder identified candidate mechanisms underlying CAR-T persistence, dysfunction and therapeutic resistance. The workflow prioritized the hypothesis that genes associated with an NK-like transition programme could be targeted to reduce CAR-T exhaustion and improve persistence. Analysis of patient scRNA-seq datasets showed enrichment of this programme in exhausted post-infusion CAR-T cells. Virtual perturbation prioritized transition-associated receptor genes, including *KLRC1*, *KLRD1* and *KLRG1*, and expert review selected *KLRC1*, encoding NKG2A, for experimental validation. *In vitro* and *in vivo* chronic-stimulation models showed that NKG2A marked activated, exhaustion-associated CD8⁺ CAR-T cells, whereas NKG2A blockade enhanced antitumour activity and persistence-associated functional readouts *in vivo*. BioPathfinder establishes a generalizable framework that transforms fragmented clinical single-cell evidence into experimentally validated therapeutic hypotheses, providing a scalable strategy for AI-guided biomedical discovery.

**HIGHLIGHTS:**

1. BioPathfinder integrates fragmented CAR-T clinical studies, patient scRNA-seq datasets and metadata into a provenance-aware evidence resource for AI-guided hypothesis discovery.
2. A multi-agent workflow generates, critiques and prioritizes diverse, falsifiable, dataset-aware mechanisms underlying CAR-T persistence, dysfunction and therapeutic resistance.
3. BioPathfinder prioritizes KLRC1/NKG2A as a therapeutic target, and experimental validation demonstrates that NKG2A blockade enhances CAR-T persistence and antitumour function.

## INTRODUCTION

Chimeric antigen receptor (CAR) T cell therapy has produced some of the most durable responses in modern cell therapy, including decade-long CD19 CAR-T persistence in patients with chronic lymphocytic leukaemia and long-term remission-associated functional programmes in pediatric acute lymphoblastic leukaemia ^1–4^ . However, durable benefit remains uneven across patients, diseases and CAR designs ^5–8^ . Clinical and single-cell studies have linked treatment failure to post-infusion cellular dynamics, regulatory CAR-T populations, inadequate persistence, tumour-intrinsic antigen escape and immunosuppressive microenvironmental states ^9–12^ . These findings suggest that patient-derived evidence contains information on the mechanisms of CAR-T persistence, dysfunction and resistance, but that such mechanisms are only interpretable in their original treatment and sampling contexts.

This evidence is difficult to use as a coherent substrate for hypothesis generation. Relevant information is distributed across primary articles, supplementary tables, GEO or other repository records, processed objects and study-specific annotations. The same biological claim can have very different implications depending on whether it is supported by infusion-product cells, early post-infusion CAR-positive cells, endogenous bystander T cells, tumour microenvironment samples, toxicity cohorts or relapse biopsies ^13^ . Therefore, the key bottleneck is not simply the availability of more CAR-T single-cell data, but the ability to connect publications, dataset accessions, sample contexts, CAR targets, clinical settings, data modalities and analysis affordances into a provenance-tracked evidence structure. Recent work has begun to place Large Language Models (LLMs) within immunotherapy discovery^14–16^. OpenIO outlined a broader AI-native immunotherapy vision built on generative AI, large-scale omics and biological foundation models ^17^ . In CAR-T research, an AI-guided study used single-cell tumour atlases, public resources and repeated LLM prioritization to nominate GPNMB as a multi-cancer antigen^18^. These studies mark important progress, but they remain distinct from an agentic, provenance-tracked system that generates mechanistic hypotheses from CAR-T-treated patient evidence.

Beyond these immunotherapy-specific efforts, related advances in AI for biology show that increasingly capable language-model and agentic workflows are beginning to support tasks in biological image analysis ^19^, computational biology ^20^, and experimental biology ^21,22^ . Retrieval-augmented language models can synthesize scientific literature with citation-backed responses ^23^, and agentic platforms have been developed for natural-language single-cell analysis ^24^, gene-set analysis ^25^, autonomous computational biology workflows and structured hypothesis generation ^21,22,26^ . These advances show that LLM-based systems can assist literature reasoning and biological data analysis. However, document retrieval or automated analysis alone does not specify which patient cohort, dataset accession, sampling time point, cell population or treatment condition supports a proposed CAR-T mechanism. For therapeutic engineering, such distinctions are essential because unsupported claims, missing dataset context and overinterpretation of computational patterns can misdirect experimental validation.

Here we present BioPathfinder, a multi-agent discovery engine for CAR-T research, which is generalizable for biomedical discovery. BioPathfinder builds a provenance-tracked paper–dataset resource from CAR-T-treated patient studies and converts it into falsifiable, dataset-aware hypotheses through a planner subagent and role-specialized LLM reviewer subagents. Applied to curated CAR-T patient single-cell evidence, the system proposes to perturb natural killer (NK)-like transition genes, which guided expert selection of *KLRC1*/NKG2A as a perturbation target ^27,28^ . We performed experiments to demonstrate that NKG2A blockade improved antitumour function and persistence, illustrating a route from structured clinical evidence to actionable CAR-T engineering hypotheses.

## RESULTS

### BioPathfinder constructs a structured evidence base from CAR-T-treated patient studies

We first asked whether dispersed CAR-T clinical evidence could be converted into a structured resource suitable for downstream hypothesis generation. BioPathfinder begins with a workflow that retrieves publications and repository records, extracts dataset accessions and evidence spans, and resolves paper–dataset links (Curator subagent). It then normalizes patient–sample contexts and CAR targets before assembling a scope-filtered frozen bundle for planning (**Fig. 1a** and **Supplementary Fig. 1a**). This design separates evidence construction from hypothesis generation, enabling subsequent subagents to operate on provenance-tracked records rather than on unstructured document retrieval.

**Figure 1.**
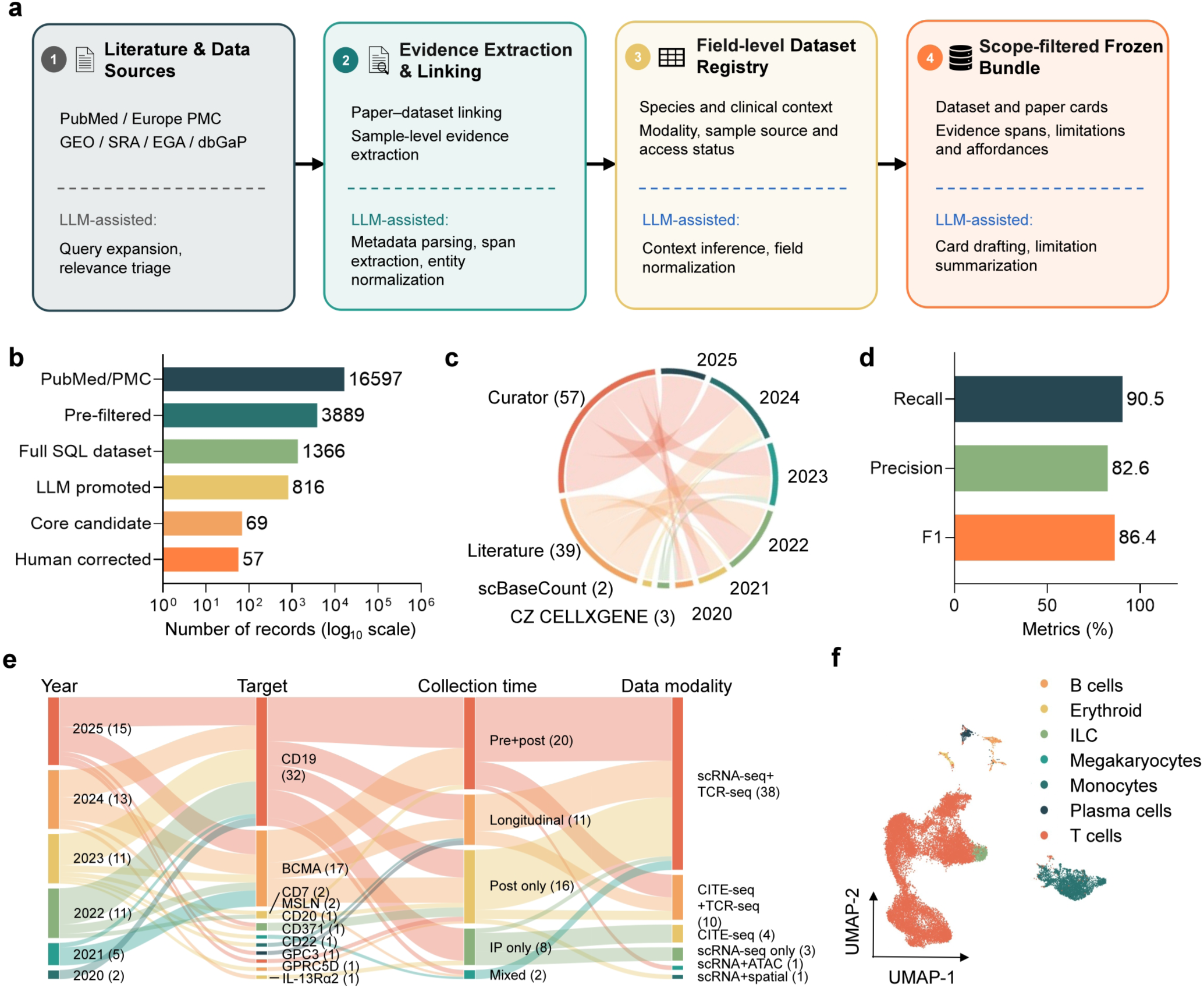
BioPathfinder constructs an expert-curated scRNA-seq resource from CAR-T cell treated patient samples. **a,** Schematic of the Curator workflow used by BioPathfinder to retrieve CAR-T cell single-cell studies, resolve literature and repository metadata, extract dataset accessions and evidence spans, adjudicate candidate records, and assemble an expert-filtered planner resource bundle. **b,** Funnel of accession-level evidence recovery from the initial PubMed/Europe PMC corpus to the final Curator core, showing the progressive reduction from broad literature retrieval to expert-approved CAR-T cell patient scRNA-seq resources. **c,** Paper-level recall comparison of Curator against external reference sources, including a recent review-derived candidate list, scBaseCount and CZ CELLxGENE coverage. **d,** Paper-level performance of the final Curator gate against the expert ground-truth set, reporting recall, precision and F1 score. **e,** Composition of the final Curator resource by publication year, CAR target antigen, sampling time point and data modality. Multi-target studies are decomposed by individual target antigen. **f,** Uniformly sampled single-cell atlas generated from Curator-supported datasets, visualized by UMAP and annotated into major cell types.

Starting from 16,597 PubMed and Europe PMC records, Curator progressively narrowed the evidence space through pre-filtering, Structured Query Language (SQL)-based dataset identification, LLM-assisted promotion and expert adjudication, returning 69 candidate publications. Manual review classified 57 as strictly eligible single-cell RNA sequencing (scRNA-seq) studies of CAR-T-treated patient with dataset-level support, 10 as patient-data-supported studies retained as supporting literature only, and two as out-of-scope records that were excluded (**Fig. 1b**). Thus, the frozen core corpus comprised 57 dataset-linked patient resources, with 10 additional supported papers available for contextual evidence. The complete inventory of curated papers and accessions is provided in **Supplementary Table 1**. Compared with external reference sources, Curator recovered a broader set of relevant papers. In the absence of a dedicated CAR-T-treated patient scRNA-seq compendium, we compared Curator with recent literature and two broad single-cell resources, scBaseCount and CZ CELLxGENE. Curator recovered 57 core studies, exceeding the 39 studies in the literature-derived list and the 2 and 3 matching studies represented in scBaseCount and CZ CELLxGENE, respectively (**Fig. 1c**) ^29–31^ . Against the expert ground-truth set, the final Curator gate achieved 90.5% recall, 82.6% precision and an F1 score of 86.4% (**Fig. 1d** and **Supplementary Fig. 1b**), indicating that the workflow prioritized high recall while retaining substantial precision.

The resulting evidence base captured key dimensions needed for CAR-T hypothesis generation, including publication year, CAR target, sampling time point and data modality (**Fig. 1e** and **Supplementary Fig. 1c**). The final resource was dominated by CD19- and BCMA-directed studies but also included less common targets such as CD7^32^, GD2^33,34^, MSLN^35^, CD22^36^, GPC3^37,38^, GPRC5D^39^ and IL-13Rα2^40^. Sampling contexts spanned pre-and post-infusion designs, longitudinal studies, post-infusion-only datasets and infusion-product-only studies. To verify that the curated records could support cell-level downstream analysis, we uniformly sampled available Curator-supported datasets and generated an annotated atlas example containing major immune and hematopoietic cell types (**Fig. 1f**) ^41,42^ . Marker-gene validation confirmed the major annotations, with T cells marked by CD3D, CD3E, CD4, CD8A and cytotoxic genes, monocyte/macrophage populations by CD14, LST1 and FCGR3A, B-lineage and plasma-cell populations by CD79A and MZB1, respectively, and erythroid cells by HBA1 and HBA2 (**Supplementary Fig. 2a–e**). Dataset-split and sample-context UMAPs further showed that the integrated atlas preserved source-dataset and treatment-context provenance, including infusion-product, pre-infusion, post-infusion and CAR-positive post-infusion sampling categories (**Supplementary Fig. 2f, g**). Together, these results establish a provenance-tracked CAR-T patient evidence base that supports reproducible multi-agent hypothesis generation.

### BioPathfinder generates and prioritizes experimentally testable CAR-T hypotheses

We next asked whether the curated CAR-T patient evidence base could be converted into a broad but reviewable hypothesis space. BioPathfinder used a planner workflow that combined curated paper findings, dataset capabilities, known gaps, an exploration question and lineage context, and then generated structured hypothesis cards containing a bounded claim, proposed analysis, confounders, stop rules and source lineage (**Fig. 2a** and **Supplementary Fig. 3a**). The planner input comprised core dataset-supported publications and additional supporting literature, and the prompt imposed fixed output fields and constraints on source use, rationale grounding, directional hypotheses, testability, dataset support and analysis readiness (Supplementary Fig. 3b). Across three repeated planner runs, the number of hypothesis–operation pairs, evidence-support pairs and unique biological targets increased steadily, indicating that the planner explored diverse mechanistic and operational hypotheses rather than repeatedly returning a narrow set of claims (**Fig. 2b**). The generated hypotheses spanned diverse candidate perturbation categories and dry-lab operation types, supporting breadth in both intervention design and computational validation strategy (**Supplementary Fig. 4a,b**). These counts were computed from schema-normalized planner outputs after exact-string and ontology-level deduplication. Hypothesis–operation pairs denote unique claim–action combinations, evidence-support pairs denote unique claim–source links, and unique biological targets denote non-redundant genes, cell states or pathways nominated for follow-up.

**Figure 2.**
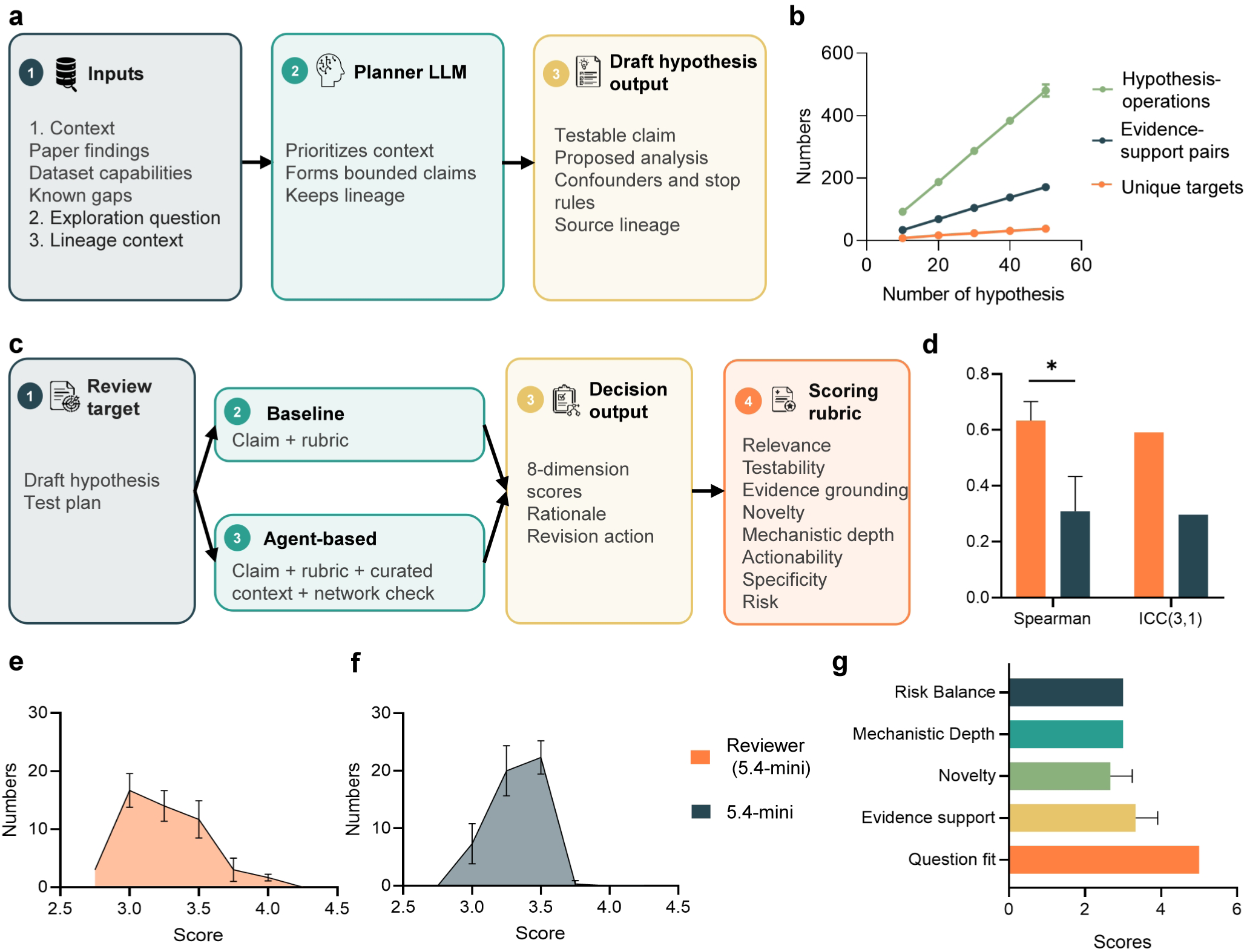
BioPathfinder generates and prioritizes experimentally testable CAR-T hypotheses. **a,** Planner workflow converting curated context, exploration questions and lineage information into structured, testable hypothesis cards. **b,** Planner output breadth across three repeated runs, measured by hypothesis–operation pairs, evidence-support pairs and unique biological targets. **c,** Reviewer workflow comparing baseline review, based on the claim and rubric, with agent-based review incorporating curated context and network checks. **d,** Repeat-reviewer reliability for hypothesis scoring using GPT-5.4-mini. The reviewer-agent setting was compared with a LLM baseline using two agreement metrics across three independent reviewer repeats: pairwise Spearman correlation and ICC(3,1). Bars show the mean of three pairwise Spearman correlations or ICC values across reviewer repeats. Error bars denote SD across repeat pairs and are descriptive only. Statistical significance was assessed for Spearman correlation by paired hypothesis-level bootstrap over 50 hypotheses using the mean Fisher-z-transformed Spearman statistic. *P = 0.016 for reviewer agent versus LLM baseline. **e,f,** Distributions of hypothesis scores assigned by the GPT-5.4-mini reviewer agent (**e**) and baseline (**f**) across the evaluated hypotheses. **g,** Dimension-level scores for the selected hypothesis, including question fit, evidence support, novelty, mechanistic depth and risk balance.

We then evaluated whether these generated hypotheses could be reproducibly prioritized. Each draft hypothesis and test plan was passed to a reviewer workflow that compared a LLM baseline, which scored the claim using the rubric alone, with an agent-based reviewer route that additionally incorporated curated context and network-check information (**Fig. 2c**). Across three repeated scoring runs over 50 hypotheses, the GPT-5.4-mini agent-based reviewer showed higher repeatability than the GPT-5.4-mini baseline, as measured by pairwise Spearman rank correlation and ICC(3,1) (**Fig. 2d**). For Spearman correlation, significance was assessed by paired hypothesis-level bootstrap using the mean Fisher-z-transformed statistic, supporting improved rank reproducibility of the agent-based reviewer route (*P = 0.016). The score distributions further showed that the agent-based reviewer produced a broader and less compressed scoring profile than the baseline, providing a less compressed score distribution for expert triage (**Fig. 2e,f**).

Finally, we asked whether the prioritized hypotheses could be converted into a biologically plausible and experimentally actionable validation plan. Human experts reviewed the top-ranked (25%) candidates for novelty, biological relevance, methodological correctness and overall scientific interest. This process selected a hypothesis: perturbing NK-like transition programme should alter exhaustion/dysfunction and cytotoxic-persistence trajectories, with chronic antigen stimulation proposed for wet-lab testing (**Table 1**)^27^ . After auditing the candidate datasets specified in the BioPathfinder hypothesis card for biological relevance, sample context and downstream analysis suitability, GSE125881 was selected as the primary patient scRNA-seq dataset for computational validation ^43^ . GSE125881 profiles isolated CD19-specific CD8^+^ CAR-T cells across infusion-product and post-infusion time points, making it well matched to testing post-infusion CAR-T state transitions and NK-like transition programme. Dimension-level scoring of the selected hypothesis showed strong question fit and evidence support, with additional support for novelty, mechanistic depth and risk-balanced usefulness (**Fig. 2g**). Together, these results show that BioPathfinder expands a structured CAR-T patient evidence base into diverse, reproducibly reviewed and expert-selectable hypotheses for computational and experimental follow-up.

**Table 1.** Selected CAR-T hypothesis after BioPathfinder and human expert nomination.

|  |  |
| --- | --- |
| **Hypothesis** | The NK-like transition observed in CAR-T cells is not merely a terminal byproduct; perturbing regulators of this transition should alter downstream exhaustion/dysfunction and cytotoxic persistence trajectories. |
| **Candidate name** | NK-like transition program |
| **Proposed relation** | Perturb |
| **Relation statement** | Perturb the NK-like transition program to redirect CAR-T cells away from dysfunction and toward durable function. |
| **Dry-lab datasets** | GSE125881 |
| **Dry-lab test** | Use GSE125881 to nominate regulators of the NK-like transition from product/post-infusion and longitudinal CAR-T state trajectories. Compute an NK-like transition program score, then test product-to-post-infusion shifts, cross-timepoint state changes, trajectory/pseudotime structure, differential expression, and marker stratification. |
| **Wet-lab test** | Perturb nominated NK-like transition regulators in primary human CAR-T cells under chronic antigen stimulation, using non-targeting controls and orthogonal rescue or alternative guides. A valid hit should preserve cytotoxic function and/or persistence while reducing dysfunction, not merely change NK-like marker frequency. |
| **Boundary** | This hypothesis can currently support candidate nomination and cross-cohort recurrence testing of NK-like transition regulators. |
| **Alternative** | NK-like states are stable terminal products; intervening upstream will not improve durable function. |

### BioPathfinder-nominated patient dataset supports perturbing NK-like transition programme in CAR-T cells

We next asked whether a BioPathfinder-prioritized hypothesis could be supported by patient-derived data and converted into perturbation candidates. In the selected dataset, unsupervised embedding separated four major T cell states: central memory-like (Tcm), effector memory-like (Tem), terminal effector memory-like (Temra), and cycling effector memory-like (Cycling Tem) populations (**Fig. 3a**) ^44^ . Across infusion product, week 2 and month 1 samples, cluster composition shifted markedly after infusion, indicating substantial remodeling of the CAR-T cell state landscape in patients (**Fig. 3b**). Differential expression analysis comparing week 2 and month 1 post-infusion samples with infusion-product cells revealed broad transcriptional rewiring after infusion (**Fig. 3c** and **Supplementary Fig. 5a**). Motivated by previous reports of a T-to-NK-like transcriptional transition, we curated a literature-derived NK-like transition programme ^28^ . To prioritize experimentally tractable candidates for downstream perturbation studies, we further defined a nested receptor-focused subset comprising genes annotated as cell-surface receptors, termed the NK-like transition receptor programme ^27^ . Gene-set enrichment analysis of week 2 and month 1 post-infusion versus infusion-product contrasts provided pathway-level context for the observed transcriptional changes (**Supplementary Fig. 5b**). In addition to broad enrichment of hallmark pathways associated with effector function and T cell dysfunction, both the literature-derived NK-like transition programme and its receptor-focused subset showed significant positive enrichment after infusion (**Fig. 3d**), indicating increased NK-like transition-associated transcriptional activity^45,46^. Together, these analyses supported the use of GSE125881 as a patient-data substrate for evaluating the BioPathfinder-nominated NK-like dysfunction hypothesis.

**Figure 3.**
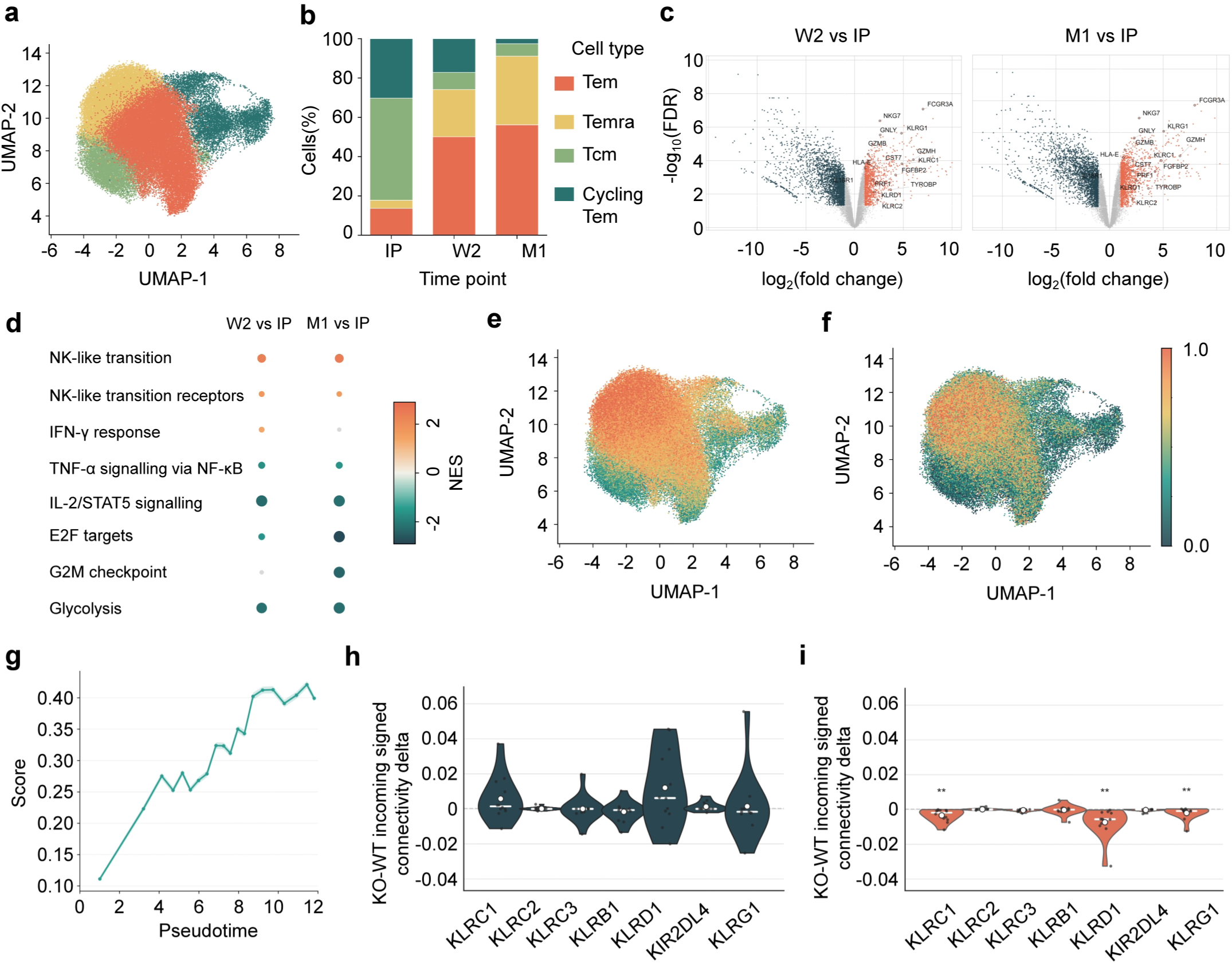
Bioinformatic validation of selected hypothesis to perturb NK-like transition programme. **a,** UMAP of clustered cells from the selected dataset. **b,** Relative cluster composition in the infusion product (IP) and post-infusion samples collected at week 2 (W2) and month 1 (M1). **c,** Differential gene expression in W2 and M1 post-infusion samples relative to IP. **d,** Dot plot of gene-set enrichment analysis for post-infusion versus IP transcriptional changes, including representative MSigDB Hallmark gene sets and literature-derived NK-like transition and NK-like transition receptor programmes. Dot colour denotes the normalized enrichment score (NES), dot size denotes −log10(FDR), and grey dots indicate FDR ≥ 0.05. **e,** UMAP visualization of NK-like transition programme scores computed by the *bioinfo_analyst* subagent. **f,** UMAP visualization of NK-like transition receptor programme scores computed by the *bioinfo_analyst* subagent. Cells in e and f are coloured by their respective programme scores. **g,** Monocle3 pseudotime analysis showing the dynamics of the NK-like transition receptor programme score along pseudotime. **h,** Knockout (KO)-induced change in incoming signed connectivity of the memory-associated programme. **i,** KO-induced change in incoming signed connectivity of the exhaustion-associated programme.

We then scored cells for NK-like transition and NK-like transition receptor programmes proposed by the analysis workflow. Module scoring showed that these programmes were preferentially enriched in defined regions of the T cell state manifold, especially in Temra-enriched regions, rather than being uniformly distributed across the manifold (**Fig. 3e,f**). Using a single principal node enriched for infusion-product Tcm cells as the trajectory root, Monocle3 ordered cells across the patient-derived T cell state manifold (**Supplementary Fig. 6a**). Along the NK-like transition branch, pseudotime increased progressively from Tcm through Tem to Temra states (**Supplementary Fig. 6b**). Along this trajectory, memory-marker activity declined, whereas cytotoxic/effector, exhaustion and NK-like transition-associated programmes increased (**Supplementary Fig. 6c**). Monocle3 trajectory analysis showed increasing NK-like transition receptor signature scores along pseudotime, consistent with a progressive acquisition of this programme during post-infusion state transition (**Fig. 3g**)^47,48^. These results support an association between post-infusion remodeling, exhausted-like T cell states and NK-like receptor programmes, while not by themselves establishing causality^49,50^.

We used virtual knockout analysis to prioritize NK-like transition receptor genes for experimental perturbation ^51^ . Candidate genes included *KLRC1*, *KLRC2*, *KLRC3*, *KLRB1*, *KLRD1*, *KIR2DL4* and *KLRG1*. Virtual perturbation of *KLRC1*, *KLRD1*, and *KLRG1* reduced the incoming signed connectivity of the exhaustion-associated programme, suggesting weakened regulatory support for the establishment or maintenance of the exhaustion-related transcriptional state (**Fig. 3h, i**). The same genes showed positive expression trends along Tcm-rooted pseudotime (**Supplementary Fig. 6d**), providing convergent temporal support for their prioritization as candidate regulators of the NK-like transition and exhaustion-associated state. We therefore treated this analysis as computational perturbation prioritization rather than causal validation. Together, scRNA-seq analysis illustrates how BioPathfinder can move from a dataset-aware hypothesis to patient-data support and a focused set of experimentally testable receptor perturbations.

### Chronic antigen stimulation enriches NKG2A-expressing CD8^+^ CAR-T cells with activated and exhausted phenotypes

Based on the predicted perturbation effects and considerations of experimental tractability, *KLRC1*, which encodes the inhibitory receptor NKG2A, was prioritized for experimental validation. This selection was supported by extensive evidence implicating NKG2A as a key NK-like inhibitory receptor associated with T cell exhaustion and dysfunctional phenotypes in CAR-T cells^52,53^. We first asked whether NKG2A marked CAR-T cells exposed to chronic tumour antigen stimulation. To model sustained tumour exposure *in vitro*, non-transduced control T cells and CD19 CAR-T cells were repeatedly co-cultured with CD19^+^ Raji human lymphoma cells and sampled across serial stimulation cycles (**Fig. 4a**) ^28^ . Flow-cytometric analysis showed that prolonged stimulation increased the fraction of NKG2A-expressing cells within the CD8⁺ CAR-T compartment (**Fig. 4b**). Quantification across the culture time course showed that NKG2A expression remained comparatively limited in non-transduced T cell controls (**Fig. 4c** and **Supplementary Fig. 7a**), whereas CAR-T cells, particularly CD8⁺ CAR-T cells, showed a progressive increase in the frequency of NKG2A⁺ cells during repeated tumour-cell stimulation (**Fig. 4d**). These data indicate that NKG2A is induced under chronic antigen exposure and is preferentially enriched in the CD8⁺ CAR-T population most directly engaged in repeated tumour-cell recognition.

**Figure 4.**
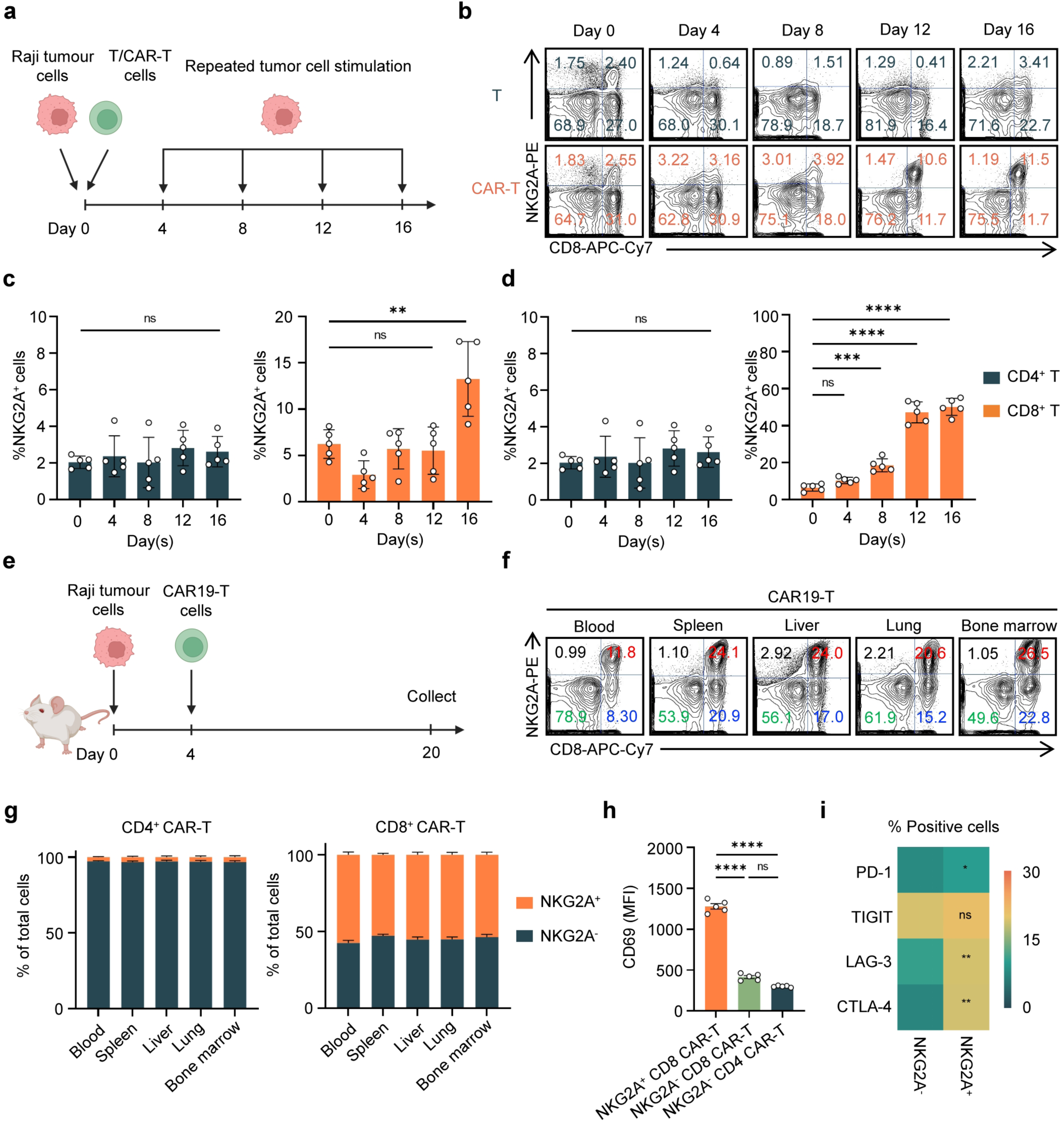
Chronic antigen stimulation enriches NKG2A-expressing CD8^+^ CAR-T cells with activated and exhausted phenotypes. **a,** Schematic of the *in vitro* repeated antigen-stimulation assay. T cells or CAR-T cells were repeatedly co-cultured with Raji tumour cells and sampled over time to assess NKG2A expression during chronic tumour-cell stimulation. **b,** Representative flow-cytometry plots showing NKG2A expression in CD8^+^ T cells and CAR-T cells after prolonged tumour-cell stimulation. Chronically stimulated CD8 CAR-T cells showed increased frequencies of NKG2A-expressing cells. **c,d,** Quantification of the frequency of NKG2A-positive cells among CD4^+^ and CD8^+^ T cells (c) or CAR-T cells (d) during repeated tumour-cell stimulation. **e,** Schematic of the *in vivo* chronic-stimulation assay. NSG mice were engrafted with Raji tumour cells and treated with CAR-T cells; CAR-T cells were collected on day 20 for phenotypic analysis after sustained *in vivo* tumour exposure. **f,** Representative flow-cytometry plots showing NKG2A expression in CAR-T cells recovered from peripheral blood, spleen, liver, lung and bone marrow. **g,** Quantification of NKG2A-positive cells among CD4^+^ and CD8^+^ T cells or CAR-T cells recovered from different mouse tissues. **h,** Activation status of NKG2A-positive and NKG2A-negative T/CAR-T cells, measured by CD69 expression. **i,** Expression of exhaustion-associated markers PD-1, TIGIT, LAG-3 and CTLA-4 in NKG2A-positive and NKG2A-negative CAR-T cells. Data are shown as mean ± s.e.m.; statistical significance was determined using one-way or two-way ANOVA with multiple-comparison correction, as appropriate. ns, not significant; *P < 0.05, **P < 0.01, ***P < 0.001, ****P < 0.0001.

We then tested whether the same phenotype emerged after sustained *in vivo* tumour exposure. NSG mice bearing Raji tumours were treated with CAR19-T cells, and CAR-T cells were collected on day 20 for phenotypic analysis after prolonged antigen encounter *in vivo* (**Fig. 4e**). NKG2A-expressing CAR-T cells were detected across multiple tissues, including peripheral blood, spleen, liver, lung and bone marrow (**Fig. 4f**). Quantification of tissue-recovered cells showed that NKG2A expression was enriched in CAR-T cells across these anatomical sites and was most prominent in the CD8⁺ CAR-T compartment compared with CD4⁺ CAR-T cells and non-transduced T cell controls (**Fig. 4g**). NKG2A⁺ CD8⁺ CAR-T cells also showed increased CD69 expression compared with NKG2A⁻ CAR-T cell subsets, linking NKG2A expression to an activation-associated phenotype after chronic stimulation (**Fig. 4h** and **Supplementary Fig. 7b**). In parallel, NKG2A⁺ CD8^+^ CAR-T cells showed increased expression of exhaustion-associated markers, with significant increases in PD-1, LAG-3 and CTLA-4 and a non-significant trend for TIGIT (**Fig. 4i** and **Supplementary Fig. 7c**). Together, these *in vitro* and *in vivo* data support NKG2A as a surface marker associated with chronic stimulation, activation and exhaustion-like phenotypes in CD8⁺ CAR-T cells.

### NKG2A blockade enhances antitumour activity of CD8^+^ CAR-T cells *in vivo*

Since NKG2A expression was enriched in chronically stimulated and exhaustion-marker-positive CD8⁺ CAR-T cells, we next tested whether blocking NKG2A could improve CAR-T cell function *in vivo*^52,53^. We used an NSG Raji-FG xenograft model in which mice were engrafted intravenously with tumour cells on day 0 and treated on day 4 with CD19-targeted CD8⁺ CAR-T cells, either alone or in combination with anti-NKG2A antibody treatment; tumour burden was monitored by bioluminescence imaging, and mice were followed for survival and terminal immune analysis (**Fig. 5a**). Representative bioluminescence images showed progressive tumour outgrowth in vehicle-treated mice, partial tumour control after CD8⁺ CAR-T cell treatment and more pronounced tumour suppression when CD8⁺ CAR-T cells were combined with anti-NKG2A blockade (**Fig. 5b**). Quantification of total bioluminescent flux confirmed reduced tumour burden over time in the combination-treatment group compared with CD8⁺ CAR-T cells alone (**Fig. 5c**). Consistent with improved tumour control, NKG2A blockade also extended survival of mice receiving CD8⁺ CAR-T cells (**Fig. 5d**).

**Figure 5.**
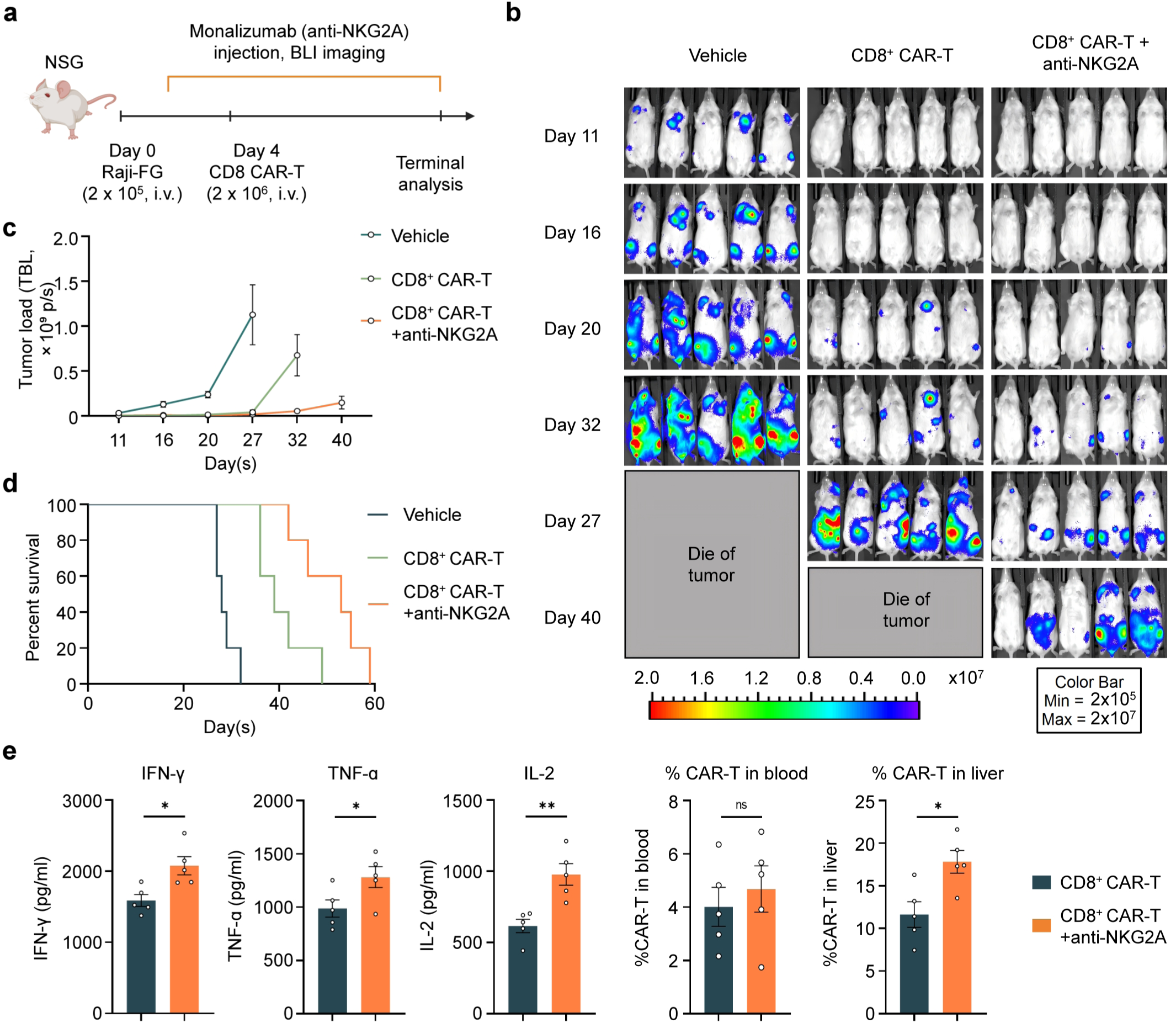
NKG2A blockade enhances CD8^+^ CAR-T cell antitumour activity in a Raji xenograft model. **a,** Schematic of the *in vivo* NKG2A-blockade experiment. NSG mice were intravenously engrafted with Raji-FG tumour cells on day 0 and treated on day 4 with CD19-targeted CD8^+^ CAR-T cells alone or CD19-targeted CD8^+^ CAR-T cells combined with anti-NKG2A antibody treatment. Tumour progression was monitored by bioluminescence imaging, and mice were followed for survival and terminal immune analysis. **b,** Representative bioluminescence images showing tumour burden over time in vehicle-treated mice, mice receiving CD8 CAR-T cells alone and mice receiving CD8^+^ CAR-T cells plus anti-NKG2A antibody. **c,** Quantification of tumour burden over time, measured as total bioluminescent flux (TBL, ×10⁹ photons s⁻ ¹). **d,** Kaplan–Meier survival curves for mice treated with vehicle, CD8^+^ CAR-T cells alone or CD8^+^ CAR-T cells plus anti-NKG2A antibody. **e,** Terminal analysis of systemic immune activation and CAR-T cell distribution, including serum IFN-γ, TNF-α and IL-2 levels, and the frequency of CAR-T cells in peripheral blood and liver. Data are shown as mean ± s.e.m.; statistical significance was determined using one-way or two-way ANOVA with multiple-comparison correction, as appropriate, or by log-rank test for survival analysis. ns, not significant; *P < 0.05, **P < 0.01, ***P < 0.001.

Terminal analysis further supported enhanced CAR-T cell activity after NKG2A blockade. Combination treatment increased systemic effector cytokine production in serum, including IFN-γ, TNF-α and IL-2, and was associated with increased CAR-T cell frequencies in liver compared with CD8⁺ CAR-T cell treatment alone (**Fig. 5e**). These findings provide functional *in vivo* evidence that NKG2A is not only a marker of chronically stimulated CD8⁺ CAR-T cells but also a therapeutically actionable inhibitory axis. Together with patient data analysis and chronic-stimulation phenotyping, these results connect the BioPathfinder prioritization path to experimental validation by connecting a computationally prioritized NK-like receptor programme to an experimentally testable intervention that improves CD8⁺ CAR-T cell antitumour activity.

## DISCUSSION

This work introduces BioPathfinder, a domain-specialized multi-agent system that converts fragmented CAR-T-treated patient evidence into experimentally actionable hypotheses. The need for such a system is underscored by various clinical therapies as exemplified in CAR-T clinical single-cell and multi-omics studies. These studies show that determinants of persistence, efficacy, toxicity and resistance are not confined to a single molecular axis, but are distributed across cellular states, tumour features, immune microenvironments, treatment contexts and molecular modalities^30,54,55^. BioPathfinder builds on, but differs from, recent retrieval-augmented and agentic systems designed for scientific literature synthesis ^23^, natural-language single-cell analysis ^26^, gene-set interpretation, autonomous biological data exploration and structured hypothesis generation ^21,22^ . Existing LLM-based and agentic approaches in CAR-T research have largely addressed predefined tasks, including antigen nomination, safety and outcome assessment, and molecular design. By contrast, to our knowledge, BioPathfinder is the first CAR-T-specific agentic workflow designed to transform provenance-tracked evidence from treated patients into diverse, falsifiable and dataset-aware mechanistic hypotheses for prioritized computational, *in vitro* and *in vivo* validation. Rather than relying on unstructured document retrieval, BioPathfinder first builds a provenance-tracked evidence base linking publications, dataset accessions, patient-sample contexts, CAR targets, modalities and analysis affordances. This structured substrate allowed planner agents to generate dataset-aware hypotheses, role-specialized LLM reviewer subagents to prioritize them, and human experts to select hypotheses for computational and experimental follow-up. By separating evidence construction, hypothesis generation, review and validation, BioPathfinder provides a reproducible route from dispersed clinical observations to testable CAR-T engineering strategies.

The biological case study highlights the utility of this framework. BioPathfinder prioritized the hypothesis that targeting genes associated with an NK-like transition programme could reduce CAR-T exhaustion and promote persistence. This hypothesis was motivated by prior evidence that CAR dysregulation can be associated with a CD8^+^ T-to-NK-like transition, and by patient studies showing that CAR-positive and CAR-negative T cells can share post-infusion differentiation trajectories into NK-like subsets^27,28^. The BioPathfinder-nominated dataset was also well matched to this question because longitudinal CD19 CAR-T profiling had previously resolved clonal kinetics and transcriptional states across infusion products and post-infusion blood samples. Patient-data analyses in this dataset supported enrichment of an NK-like transition-associated programme in exhausted CAR-T states and nominated a focused set of receptor genes for perturbation. Subsequent expert selection and experimental validation identified *KLRC1*, encoding the inhibitory receptor NKG2A, as a functional modulator of CAR-T dysfunction. This target choice is consistent with prior patient single-cell work in which *KLRC1* was observed within post-infusion NK-like CAR-T or NK-like T cell signatures. Importantly, however, the functional perturbation result is established by the present study: *KLRC1*/NKG2A is associated with sustained antigen stimulation and exhaustion phenotypes, and its blockade improved CAR-T antitumoural function and persistence *in vivo*. These findings suggest that NK-like inhibitory receptor programmes represent a targetable axis of post-infusion CAR-T dysfunction. More broadly, such molecular targets could be integrated with emerging biomaterial-enabled strategies for *ex vivo* cell manufacturing, targeted delivery and *in vivo* immune-cell modulation to further improve the potency, persistence and accessibility of next-generation CAR-T therapies^8^.

Our results also clarify the role of multi-agent systems in biomedical discovery. Recent biological AI agents and AI co-scientist systems have demonstrated that language-model-based agents can support literature synthesis, biological database verification, computational analysis and hypothesis generation ^25,56^ . BioPathfinder extends this direction in a therapeutic context by emphasizing provenance, dataset scope, evidence boundaries and expert-mediated claim control. It did not replace expert judgment or autonomously establish causal mechanisms. Instead, it expanded the hypothesis search space, organized evidence boundaries, exposed confounders and enabled structured prioritization before wet-lab testing. This human-guided design is important for therapeutic research, where unsupported claims, incomplete dataset context and overinterpretation of computational analyses can misdirect experiments. The role-specialized LLM reviewer subagents further provided complementary assessments of biological plausibility, dataset suitability, feasibility, risk and claim safety, aiding to convert generated hypotheses into experimentally responsible proposals.

While BioPathfinder offers the advantage of prioritizing clinically relevant, patient-derived single-cell and multi-omics evidence that captures treatment context, cellular states and response-associated biology, several limitations remain to be addressed. First, the current evidence base is intentionally restricted to studies of patients treated with CAR-T cells, which may narrow the mechanistic search space by excluding potentially informative evidence from model systems, manufacturing studies and unpublished datasets ^57,58^ . Additionally, the quality of the generated hypotheses depends on the completeness and accuracy of the extracted metadata and evidence spans. In particular, unresolved paper–dataset relationships and incomplete sample annotations remain challenging to identify automatically and may limit the comprehensiveness of the resulting hypotheses. The bioinformatic analyses provide support and prioritization, but causal interpretation still requires perturbation experiments. BioPathfinder is not yet a fully automatic discovery system, human input remains necessary for defining analysis scope, resolving ambiguous evidence links, selecting perturbation targets, interpreting computational outputs and making experimental decisions. Future implementations could more tightly couple hypothesis cards to executable patient-data analyses, automated visualization and manuscript-assembly workflows, thereby reducing manual intervention while preserving expert control over biological claims. Finally, although *KLRC1*/NKG2A emerged as a validated target in this study, broader prospective testing across additional CAR constructs, tumour models and patient contexts will be needed to determine the generality of NK-like receptor perturbation as a CAR-T engineering strategy^33,37,40,59^.

In summary, BioPathfinder demonstrates that structured clinical evidence and multi-agent reasoning can be combined to generate experimentally actionable hypotheses in cell therapy. By linking evidence curation, hypothesis generation, expert-style review and experimental validation, this framework offers a generalizable model for domain-specific AI-assisted discovery in translational biomedicine.

## METHODS

### BioPathfinder architecture and scope

BioPathfinder was implemented as a modular multi-agent workflow for CAR-T single-cell source curation, hypothesis generation, hypothesis review and downstream bioinformatics planning. The workflow was comprised of four linked agentic components. The Curator component performed source extraction and database construction by converting literature and official repository records into structured paper, dataset and source-artifact records. Planner used the frozen Curator-derived artifacts and a prespecified exploration question to generate bounded draft hypotheses with machine-readable lineage and reviewer-facing claim-plan views. Reviewer evaluated the same standardized hypothesis view through LLM baseline and agent-assisted routes. Bioinfo Analyst translated reviewer-selected hypotheses into executable, human-gated bioinformatics analysis plans.

This architecture separated source curation, hypothesis generation, merit review and downstream analysis planning into explicit stages. Upstream source artifacts were frozen before hypothesis generation, allowing Planner and Reviewer to operate on a defined, traceable evidence substrate. The reviewer-facing claim-plan view retained the hypothesis identifier, title, core claim, proposed dry-lab and wet-lab tests, claim boundary and strongest alternative, while omitting generic rationale and other presentation-prone fields from scored review. Runtime identifiers, schema manifests, model settings and run-level provenance are reported in the reproducibility materials.

### Curator evidence extraction and database construction

Curator was implemented as the source-curation and database-construction component of BioPathfinder. It began with publication-level candidate discovery rather than accession-first retrieval. For the manuscript freeze, Europe PMC was queried for records dated from 1 January 2017 to 4 May 2026 using CAR-T terms crossed with single-cell or multiomic profiling terms. This search returned 16,597 publication records; a title- and abstract-level prefilter retained 3,889 records with CAR-T construct, product, clinical-study or initial inclusion signals for resolver-based identifier recovery and source grounding.

For retained publications, resolver and grounding services recovered stable publication identifiers, repository identifiers and source contexts, including data-availability statements, methods text, supplementary links, repository pages and accession-centred text windows. These steps produced source-grounded paper-accession candidates. Candidate accessions were first treated as repository entry points. Each accession-level candidate was represented by a source-grounded packet containing publication metadata, accession-neighbouring source windows and available repository metadata. The grounded accession-level candidates then entered an LLM-assisted adjudication layer. LLM-assisted classifiers were used to assess eligibility and normalize fields relevant to downstream use, including CAR-T context, study context, species or sample origin, clinical workflow, single-cell modality, access level, analysis readiness, source role and limitations. LLM outputs were treated as adjudication signals and structured projections, whereas repository metadata and source windows remained the authoritative records for dataset facts. Missing or unresolved fields were retained as unknown. Curator exported the resulting records into relational database tables and typed artifacts linking papers, accessions, dataset-source records, dataset records, sample anchors and source provenance. The rule-based calibration step produced a clinical CAR-T single-cell core of 69 accession-level records. After source-context construction and expert calibration, these were consolidated into 57 curated accession-source records for downstream planning and review. Planner and Reviewer then used the retained paper records, source windows, accession-source records and accession indexes for hypothesis generation and review.

### Planner hypothesis generation

Planner generated hypotheses from an exploration question and selected upstream artifacts rather than from unrestricted retrieval. The frozen manuscript run used paper and dataset selection, a structured draft-hypothesis generator, a frontier-discovery planning track and structure-preserving post-processing. The reference evaluation run requested up to ten hypotheses for each exploration question and emitted 50 schema-valid normalized hypotheses. Planner retained two views of each generated hypothesis. A machine-rich record preserved lineage, selected artifacts, source-resolution fields, domain priors and dry-lab or wet-lab planning structure. The reviewer-facing view retained the hypothesis identifier, title, core hypothesis, candidate intervention or state, dry-lab test, wet-lab test, claim boundary and strongest alternative. Generic rationale text and self-certifying planning fields were omitted from this scored view to reduce presentation-driven bias. Reviewer scoring used this claim-plan view to compare hypotheses on a common textual basis, while the machine-rich record preserved provenance for source resolution and audit. Generation outputs were validated against local schemas, and only valid structured outputs advanced to downstream review.

### Reviewer routing and hypothesis-merit reproducibility evaluation

The reviewer-facing unit was the bias-controlled draft-hypothesis view. For each hypothesis, the pipeline produced baseline, and agent-assisted review packets that shared the same scored target. The LLM baseline route measured intrinsic hypothesis quality from the exploration question and claim-plan text. The agent-assisted route combined local corpus inspection from Curator result, with live literature search to assess novelty, capability requirements and claim boundaries. Each route emitted the same structured hypothesis-merit scorecard. Primary scientific-merit dimensions were question fit, claim clarity, evidence-bridge support, novelty, mechanistic depth, decision value and risk-balanced usefulness; secondary readiness dimensions were dry-lab readiness, wet-lab readiness and boundary quality. For cross-route reliability, the shared primary score averaged dimensions available in all routes and reported source-dependent evidence-bridge support separately. Reviewer reproducibility was evaluated on 50 reviewer-facing hypotheses by repeating scoring three times for each evaluated model-route condition. Rank stability and agreement across repeats were summarized using pairwise Spearman correlation, Kendall tau-b, top-ranked overlap and intraclass correlation. Reviewer outputs were used to prioritize hypotheses for author review and validation planning. To assess the reproducibility of Planner output, the Planner generated 50 research questions in each of three independent run sequences. Supplementary Fig. 4 presents additional analyses in Fig. 2b, rather than results from a separate experiment. For reviewer evaluation, each reviewer type independently scored 50 Planner-generated questions in each of three repeated runs; the resulting scores were used for the analyses shown in Fig. 2d–f.

### Bioinfo Analyst tool registry and human-gated execution

Bioinfo Analyst was implemented as a controlled local analysis orchestration layer. The registry was limited to local tools, accepted only declared parameters and required user-supplied gene programmes for programme-based analyses. Registered tools included quality control and preprocessing, scVI clustering, Seurat RPCA integration, cluster marker testing, cluster composition, gene-expression UMAP plotting, programme scoring, pseudobulk generation, DEG/GSEA, Monocle3 or CellRank-compatible trajectory analysis, programme-pseudotime analysis and virtual knockout. Reviewer-accepted targets were converted into structured bioinformatics task records only when the reviewer decision allowed analysis. Tasks remained in dry-run mode until required local input files, parameters and gene programmes were supplied. Execution plans held incomplete tasks in a planning state and emitted executable command arguments for input-complete tasks. Bioinformatics result records described execution status and output artifacts, and result interpretation was routed back to reviewer or human review. Virtual knockout was reserved for explicitly selected perturbation analyses.

### Single-cell QC and preprocessing

Single-cell preprocessing was performed with Scanpy, AnnData, NumPy, pandas, SciPy and Scrublet. Raw count matrices were obtained from AnnData objects or 10x Genomics-formatted matrices. When available, raw counts stored in layers[“counts”] were used as the input for quality control. Cells were filtered using a minimum gene count threshold of 200 and a mitochondrial read fraction threshold of 15% unless otherwise specified. Ambient-like cells were flagged using a predefined ambient gene list, including haemoglobin, platelet and myeloid genes, followed by a quantile-based ambient fraction cutoff. Doublet detection was performed with Scrublet using an expected doublet rate of 0.06.

Actual GSE125881 collection times were CLL-1: IP, d21, d38, d112; CLL-2: IP, d12, d29, d83; NHL-6: IP, d12, d29, d102; and NHL-7: IP, d12, d28, d89; for the pooled composition labels, IP denotes the infusion product, W2 includes d12 only, M1 combines d21/d28/d29/d38, and the late d83/d89/d102/d112 samples are excluded.

After quality control, raw counts were retained in layers[“counts”] for downstream modelling. Highly variable genes were selected with Scanpy using the Seurat v3 method when available, with fallback to Cell Ranger-style highly variable gene selection after normalization. The default number of highly variable genes was 3,000 unless otherwise specified. TCR/BCR variable-region genes and selected sex-linked genes were excluded from highly variable gene selection using predefined gene symbols and gene-prefix filters. Processed all-gene and highly variable gene AnnData objects were retained for downstream analyses.

### scVI clustering and CellTypist annotation

Latent integration and clustering were performed with scVI-tools and Scanpy. AnnData objects were configured to use layers[“counts”] as the count input. The default batch key was sample_id. Optional categorical and continuous covariates were included only when present in the cell metadata. The scVI model used a 30-dimensional latent space, two neural network layers, 128 hidden units, a dropout rate of 0.1 and a negative binomial likelihood by default. Model training used GPU acceleration when available and otherwise used CPU execution.

The scVI latent representation was stored in obsm[“X_scVI”]. Nearest-neighbour graphs were computed from this latent representation, followed by UMAP visualization and Leiden clustering. Default Leiden resolutions were 0.2, 0.3, 0.5, 0.8, 1.0, 1.2, 1.5 and 2.0. Cell-type annotation was performed with CellTypist on count-derived log-normalized expression using the specified immune reference model. Immune_All_Low.pkl was used as the default model. Per-cell CellTypist labels and cluster-majority CellTypist summaries were used for downstream interpretation.

### Cluster marker analysis and composition analysis

Cluster marker genes were identified with Scanpy rank_genes_groups. The default statistical test was the Wilcoxon rank-sum test. Clusters containing fewer than 100 cells were excluded from marker testing by default. Marker outputs included gene rank, test score, nominal P value, adjusted P value and log fold change. Cluster composition was calculated by grouping cells according to cluster labels and available sample-level metadata, including sample context, time point or clinical group when available.

### Pseudobulk differential expression and GSEA

Pseudobulk count matrices were generated by summing raw counts across cells within each sample and cell-type or cluster group. Groups containing fewer than 20 cells were excluded by default. Differential expression and pathway enrichment were performed with an R backend. Pseudobulk differential expression used edgeR quasi-likelihood generalized linear models with TMM normalization. Paired analyses used a ∼ patient + group design, whereas unpaired group comparisons used a ∼ group design. Genes were ranked for enrichment analysis using sign(logFC) × -log10(PValue). Gene set enrichment analysis was performed with fgsea using MSigDB Hallmark gene sets and optional user-supplied gene programmes.

The NK-like transition-associated gene programme (NK-like transition) comprised *NKG7, GNLY, PRF1, GZMB, GZMH, CST7, FGFBP2, KLRC1, KLRC2, KLRC3, KLRB1, KLRD1, KLRG1, KIR2DL4, FCGR3A, TYROBP, IL18R1* and *HLA-E*, whereas the receptor-focused programme (NK-like transition receptors) comprised *KLRC1, KLRC2, KLRC3, KLRB1, KLRD1, KIR2DL4* and *KLRG1*, representing an NK-receptor-associated subset of the broader programme; both gene sets were curated on the basis of NK-like CAR-T transition signatures reported by Louie et al. and Good et al^27,28^.

### Pseudotime analysis

Pseudotime analysis was performed with Monocle3 through an R backend. Input AnnData objects were required to contain raw counts in layers[“counts”], UMAP coordinates and a cluster annotation. Root cells were selected either from a user-specified root cluster or according to high or low expression of a supplied gene set. Count matrices, cell metadata, gene annotations, UMAP coordinates and root-cell assignments were passed to Monocle3. Pseudotime values were then imported back into AnnData for downstream analysis of gene-programme dynamics. CellRank-compatible arguments were retained for interface compatibility but were not used in the current implementation.

### Virtual knockout

Virtual knockout analysis was performed with scTenifoldKnk through an R backend. Input AnnData objects were converted to 10x Genomics-style matrix format when required by the backend. The knockout gene, cell-label pattern and gene-programme definition were specified by the user for each analysis; no default knockout gene was used. For each candidate gene, scTenifoldKnk inferred perturbation effects and summarized programme-level changes across selected cell states. Default parameters included a maximum of 800 genes, four network replicates, 300 cells per run, tensor decomposition rank of 2, two CPU cores. Virtual knockout results were interpreted as computational perturbation prioritization rather than causal validation.

### Mice

This study utilized the NOD.Cg-Prkdc^SCID^Il2rg^tm1Wjl^/SzJ (NOD/SCID/IL-2Rγ^−/-^, NSG) mice housed by the animal facilities of the University of California, Los Angeles (UCLA). The mice used in experiments aged between 6-10 weeks, unless otherwise specified. Both male and female mice were included in the *in vivo* assays, as no sex-based differences were observed in tumour growth or therapeutic efficacy in this model. All protocols (ARC-2013-054) of animal experiments were reviewed and approved by the Institutional Animal Care and Use Committee (IACUC) of UCLA. All mice were housed under specific pathogen-free conditions, and all experimental procedures were conducted in accordance with the regulations established by the Division of Laboratory Animal Medicine (DLAM) at UCLA.

### Cell lines

Human Burkitt’s lymphoma cell line Raji and human embryonic kidney (HEK) cell line 293T were obtained from the American Type Culture Collection (ATCC). To generate stable tumour cell lines overexpressing the firefly luciferase and enhanced green fluorescence protein dual-reporters (FG), the parental tumour cell line was transduced with lentiviral vectors encoding the intended gene(s). 72 h post lentivector transduction, tumour cells were subjected to flow cytometry sorting to isolate gene-engineered cells for making stable cell lines. Raji-FG tumour cell line was generated for this study. All tumour cell lines used in this study were tested mycoplasma-free and frequently STR verified during experiments.

### Human peripheral blood mononuclear cells (PBMCs)

Healthy donor PBMCs were obtained from the UCLA/CFAR Virology Core Laboratory, without identification information under federal and state regulations. PBMCs were cryopreserved in Cryostor CS10 medium using CoolCell and were frozen in liquid nitrogen for storage and to supply all experiments.

Recombinant human IL-2 (cat. no. 200-02) was purchased from PeproTech. Monalizumab (cat. no. HY-P99032) was purchased from MCE. Fetal bovine serum (FBS) (cat. no. F2442), and β-mercaptoethanol (β-ME) (cat. no. M3148) were purchased from Sigma. Penicillin-streptomycin-glutamine (P/S/G) (cat. no. 10378016), MEM nonessential amino acids (NEAA) (cat. no. 11140050), HEPES buffer solution (cat. no. 15630080), and sodium pyruvate (cat. no. 11360070) were purchased from Gibco. Normocin (cat. no. ant-nr-05) was purchased from InvivoGen. The RPMI 1640 cell culture medium (cat. no. 11875093) and the DMEM cell culture medium (cat. no. 11965092) were purchased from Thermo Fisher Scientific. The CryoStor Cell Cryopreservation Media CS10 (cat. no. C2999) was purchased from MilliporeSigma.

The C10 medium was made of RPMI 1640 cell culture medium supplemented with FBS (10% v/v), P/S/G (1% v/v), NEAA (1% v/v), HEPES (10 mM), sodium pyruvate (1 mM), β-ME (50 μM), and Normocin (100 μg/ml). The C10 medium was used to culture human CAR-T cells. The D10 medium was made of DMEM supplemented with FBS (10% v/v), P/S/G (1% v/v), and Normocin (100 μg/ml). The D10 medium was used to culture HEK 293T cells. The R10 medium was made of RPMI 1640 supplemented with FBS (10% v/v), P/S/G (1% v/v), and Normocin (100 μg/ml). The R10 medium was utilized to culture human tumour cells such as Raji cells.

### Lentiviral vectors

All lentiviral vectors used in this study were constructed from a parental vector pMNDW. ^60,61^ The 2A sequence derived from porcine teschovirus-1 (P2A) was used to link the inserted genes to achieve co-expression. The Lenti/FG vector was constructed by inserting a synthetic bicistronic gene encoding Fluc-P2A-EGFP into the pMNDW. ^60,62^ The Lenti/CAR19 vector was constructed by inserting a synthetic gene encoding human CD19-targeting CAR, consisting of an anti-CD19 scFv, CD8 hinge, CD8 transmembrane domain, and CD28 and CD3ζ signaling domains, into the pMNDW. ^63^ The synthetic gene fragments were obtained from GenScript and IDT. Lentiviruses were produced using HEK 293T cells, following a standard transfection protocol using the Trans-IT-Lenti Transfection Reagent (Mirus Bio) and a centrifugation concentration protocol using the Amicon Ultra Centrifugal Filter Units, according to the manufacturer’s instructions (MilliporeSigma).

### Antibodies and flow cytometry

Fluorochrome-conjugated antibodies specific to human CD45 (Clone HI30, cat. no. 304026, 1:500 dilution), CD3 (Clone HIT3a, cat. no. 300329, 1:500 dilution), CD4 (Clone OKT4, cat. no. 317414, 1:400 dilution), CD8 (Clone SK1, cat. no. 344714, 1:300 dilution), CD69 (Clone FN50, cat. no. 310906, 1:50 dilution), and NKG2A (Clone S19004C, cat. no. 375103, 1:50 dilution) were purchased from BioLegend. Fixable Viability Dye eFluor506 (e506, cat. no. 65-0866-14, 1:500 dilution) was purchased from Affymetrix eBioscience; mouse Fc Block (antimouse CD16/32, cat. no. 553142, 1:50 dilution; RRID: AB_394656) was purchased from BD Biosciences. A goat anti-mouse IgG F(ab’)2 secondary antibody (HRP-conjugated, 1:50, cat. no. 31436, RRID: 228313) was purchased from Thermo Fisher Scientific. All flow cytometry staining was performed following standard protocols, as well as specific instructions provided by the manufacturer of a particular antibody. Stained cells were analysed using a MACSQuant Analyser 10 flow cytometer (Miltenyi Biotech), following the manufacturer’s instructions. FlowJo software v.9 (BD Biosciences) was used for data analysis.

### Enzyme-linked immunosorbent cytokine assays (ELISA)

For human IFN-γ, TNF-α and IL-2 detection, capture and biotinylated antibody pairs were purchased from BD Biosciences, streptavidin–HRP conjugate was purchased from Invitrogen, human cytokine standards were purchased from eBioscience, and tetramethylbenzidine (TMB) substrate was purchased from KPL. The samples were analysed for absorbance at 450 nm using an Infinite M1000 microplate reader (Tecan).

### Generation of human CD19-targeting conventional CAR-T cells

Non-treated tissue culture 24-well or 12-well plates (Corning) were coated with Ultra-LEAF^TM^ Purified Anti-Human CD3 Antibody (Clone OKT3; BioLegend) at 1 μg/ml (500 μl/well), at room temperature for 2 hours or alternatively, overnight at 4 °C. Healthy donor PBMCs were resuspended in the C10 medium supplemented with 1 μg/ml Ultra-LEAF^TM^ Purified Anti-Human CD28 Antibody (Clone CD28.2, BioLegend) and 30 ng/ml IL-2, followed by seeding in the pre-coated plates at 1 × 10^6^ cells/ml (1 ml/well). On day 2, cells were transduced with Lenti/CAR19 viruses for 24 hours. The resulting CAR-T cells were expanded for about 2 weeks in C10 medium supplemented with human IL-2, and cryopreserved for future use, following established protocols.^64,65^

### *In vivo* bioluminescence imaging (BLI)

BLI was conducted using the Spectral Advanced Molecular Imaging (AMI) HTX system (Spectral Instrument Imaging). Live animal images were acquired 5 min after intraperitoneal (i.p.) injection of D-luciferin to capture whole-body bioluminescence signals. To monitor tumour cells, 1 mg of D-luciferin was administered per mouse. To monitor therapeutic cells, 3 mg of D-luciferin was administered per mouse. Imaging data were processed and analysed using AURA imaging software (Spectral Instrument Imaging, version 3.2.0).

### *In vitro* repeated antigen-stimulation assay of CAR-T cells

Experimental design is shown in Fig. 4a. CAR-T cells or untransduced T cells were co-cultured with Raji tumour cells. To model chronic antigen exposure, fresh Raji tumour cells were added every 4 days for a total duration of 16 days. At the indicated time points, cells were collected for flow-cytometric analysis of NKG2A expression on CD4^+^ and CD8^+^ T cell populations.

### *In vivo* chronic stimulation assay of CAR-T cells

On Day 0, NSG mice were intravenously inoculated with Raji tumour cells (2 × 10^5^ cells per mouse). On Day 4, mice received CAR-T cells by intravenous injection. Mice were maintained for 20 days to allow sustained *in vivo* tumour exposure and chronic antigen stimulation. Tumour burden was monitored by BLI throughout the study. On Day 20, mice were euthanized and CAR-T cells were isolated from peripheral blood, spleen, liver, lung, and bone marrow for phenotypic analysis. NKG2A expression was assessed by flow cytometry on CD4^+^ and CD8^+^ CAR-T cell subsets. Activation and exhaustion markers were quantified in NKG2A^+^ and NKG2A^−^ CAR-T cell populations.

### *In vivo* NKG2A immune checkpoint blockade study of CD8^+^ CAR-T cells

Experimental design is shown in Fig. 5a. On Day 0, NSG mice were intravenously inoculated with Raji-FG cells (2 × 10^5^ cells per mouse). On Day 4, mice received one of the following intravenous treatments: vehicle control (100 μL PBS per mouse), CD8^+^ CAR-T cells (2 × 10^6^ cells in 100 μL PBS per mouse), or a combination of CD8^+^ CAR-T cells (2 × 10^6^ cells in 100 μL PBS per mouse) with anti-NKG2A antibody (Monalizumab, 10 mg/kg). For groups receiving anti-NKG2A treatment, the antibody was administered every 7 days throughout the duration of the experiment. Mice were monitored for survival, and tumour burden was assessed using BLI. At the end of the experiment, mice were euthanized, and their tissues and serum were collected for CAR-T cell distribution and functional analysis by flow cytometry and ELISA.

### Statistics

Student’s two-tailed t-test was used for pairwise comparisons. Ordinary one-way analysis of variance (ANOVA) followed by Tukey’s or Dunnett’s multiple comparisons test was used for multiple comparisons. A log rank (Mantel-Cox) test adjusted for multiple comparisons was used for Meier survival curves analysis. Data are presented as the mean ± s.e.m., unless otherwise indicated. In all figures and figure legends, n represents the number of samples or animals used in the indicated experiments. A P value of less than 0.05 was considered significant and NS denotes not significant.

### Reproducibility

All agent-facing stages wrote structured metadata sufficient to audit the run context. The manuscript workflow stored an upstream manifest, stage-level outputs, selected artifact references, generation outputs, normalized hypotheses, reviewer-facing claim-plan views, route-specific review packets, prompts, raw responses, scorecards, failure files and calibration summaries. The frozen upstream inputs included structured dataset-source records, dataset cards, paper source-window artifacts and dataset-card indexes. Planner LLM runs recorded runtime schemas, model configuration, prompt and schema hashes, request hashes, artifact origin, validation status and output locations. Hypothesis-merit reviewer runs recorded route, model identifier, repeat label, prompt, raw response, normalized scorecard, failures and summary statistics. Agent-assisted reviewer runs additionally stored work orders, raw agent outputs, thread metadata and search-trace records. Bioinfo execution plans stored input-file references, accepted parameters, gene-programme references, command previews, blocking reasons and output locations. Random seeds were fixed where the underlying tool exposed a seed. Frozen upstream artifacts, schema validation and structured-output checks made the run auditable. Detailed file paths, internal schema identifiers and software commit information were retained in the code and data release rather than in the main manuscript Methods. Planner and reviewer outputs were used to prioritize hypotheses for downstream computational and experimental validation.

## Supporting information

Supplementary Table 1_Complete inventory of curated papers and accessions

## Data availability

The data supporting the findings of this study are available within the manuscript and its Supplementary Information. The publicly available single-cell RNA-sequencing dataset analysed in this study is available from the Gene Expression Omnibus under accession number GSE125881. Additional data is available from the corresponding author upon reasonable request.

## Code availability

Source code and analysis scripts are available from the BioPathfinder GitHub repository at https://github.com/TeammateDownloadGenshin/BioPathfinder. The version associated with this study is archived as release v1.0.0-rc3 at https://github.com/TeammateDownloadGenshin/BioPathfinder/releases/tag/v1.0.0-rc3.

## Acknowledgements

We thank the UCLA/CFAR Virology Core Laboratory for providing healthy-donor peripheral blood mononuclear cells and the UCLA Division of Laboratory Animal Medicine for animal care and support.

## Author contributions

S.W., Y.-R.L. and Q.W. designed the experiments, analysed the data and wrote the manuscript. S.W., Q.W., Y.Y., Z.C., H.L., H.D., X.H. and Y.Z. developed the software. S.W., Y.Y., Z.C. and H.N. performed the bioinformatic analyses. S.W., Y.-R.L., Q.W., Y.Y., X.S., H.L., H.N., Z.C., Y.C.Z., B.Z., H.D., J.S., S.P., Y.C.Z. and

X.H. contributed to the *in vitro* and *in vivo* experiments and data curation. Y.-R.L., D.L. and S.L. conceived and supervised the study, interpreted the results and revised the manuscript. All authors reviewed and approved the final manuscript.

