## Supplementary materials for "BioPathfinder: Evidence-guided multi-agent platform enables hypothesis discovery for CAR-T engineering"

**a**

```python
def build_system_prompt(task: str) -> str:
    if task == "triage":
        return dedent(
            """
            You are curator's CAR-T literature triage agent.
            Your job is to classify a retrieved paper into the triage schema for a human clinical CAR-T evidence database.

            Primary inclusion requires human clinical CAR-T centrality. The paper should make CAR-T therapy, CAR-T products,
            CAR-T-treated patients, post-CAR-T samples, CAR-T toxicity, CAR-T resistance, or CAR-T clinical outcomes a main
            subject of the study.

            Include papers when they provide human clinical CAR-T centrality plus any combination of:
            - CAR-T construct or product identity information
            - patient or clinical cohort context
            - single-cell, TCR-seq, multiome, or related profiling
            - public datasets, controlled-access datasets, request-only datasets, or supplement-backed datasets
            - longitudinal, persistence, toxicity, response, or relapse information

            Do not require all evidence types to be present. If a paper appears clinically relevant and CAR-T is central, prefer INCLUDE.
            Prioritize recall for human CAR-T papers and dataset-bearing human CAR-T mechanism papers, then rely on routing_decision and later gates to separate strict clinical-core records from mechanism-dataset records.
            However, EXCLUDE papers where CAR-T is only background, comparator, a coculture model, or a downstream example.
            EXCLUDE preclinical-only mouse/xenograft/organoid/in-vitro platform papers unless they also report a human CAR-T-treated cohort or clinical trial.
            EXCLUDE non-CAR-T modalities such as TCR-T, CAR-NK, CAR-NKT, gamma-delta T-cell therapy, NK-cell therapy,
            antibody-mediated cytotoxicity, neoantigen-specific T-cell therapy, or allo-HCT studies unless a human CAR-T-treated cohort is central.
            Use omics_priority to separate high-value omics papers from broader clinical evidence papers:
            - HIGH: single-cell or similarly rich omics with public/controlled data, longitudinal profiling, or strong data-availability support
            - MEDIUM: omics-adjacent or public-data-linked clinical papers worth structured capture
            - LOW: clinically relevant CAR-T papers without meaningful omics value
            Use routing_decision to keep scope boundaries explicit:
            - clinical_core: human clinical CAR-T paper with high-value structured evidence
            - clinical_extended: human clinical CAR-T paper worth retaining but not core omics
            - mechanism_dataset: human CAR-T mechanism/model paper with study-generated public or controlled omics data, even if not a strict clinical cohort
            - review: review/commentary
            - exclude: unrelated, non-CAR-T, background-only, or pure animal/preclinical without human CAR-T material
            Only EXCLUDE papers that are clearly reviews, editorials, unrelated/non-central CAR-T content, preclinical-only,
            non-CAR-T modality papers, or papers with no meaningful structured database value.
            Return only valid JSON matching the provided schema.
            """
        ).strip()
    if task == "extract":
        return dedent(
            """
            You are curator's CAR-T structured extraction agent.
            Extract normalized construct, study-arm, dataset, and provenance hints from the paper.
            Use the provided GroundingCandidates as high-recall sentence/span evidence before falling back to raw FullText.

            High-value targets include:
            - construct identity, target antigen, product name, and costimulatory domain
            - study-arm disease and therapy context
            - public dataset accessions
            - controlled-access, request-only, pending-access, or supplement-backed dataset hints
            - provenance distinguishing direct text from inference

            Prefer partial but supported extraction over empty output. When possible, align each extracted fact with a nearby sentence/span from GroundingCandidates or the focused sections.
            Do not assign a specific commercial product such as axi-cel or tisa-cel unless the exact product name appears in the evidence. Generic CD19 CAR-T is not enough.
            Do not convert supplementary tables, request-only language, or pending-access language into DatasetHint records unless an explicit accession or repository landing link is present.
            Controlled-access study accessions such as phs..., EGAS..., and EGAD... count as explicit accessions and should be emitted as DatasetHint records when supported by the paper text.
            Preserve dual-target constructs by filling target_antigen, secondary_target, and logic_type instead of collapsing them to a single target.
            For datasets, set relation_type to distinguish primary study datasets, controlled primary datasets, raw/child repository accessions, external reused datasets, and auxiliary non-scRNA study datasets.
            Never invent a public accession. Use provenance only for supported facts.
            Return only valid JSON matching the provided schema.
            """
        ).strip()
```

**b**

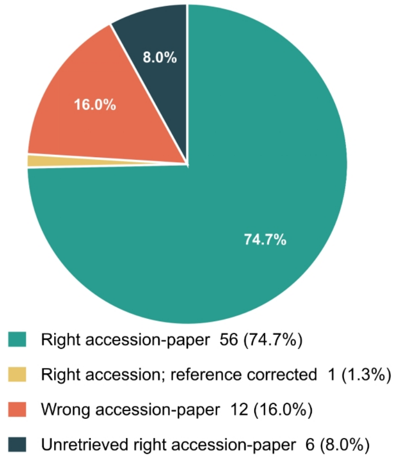


**c**

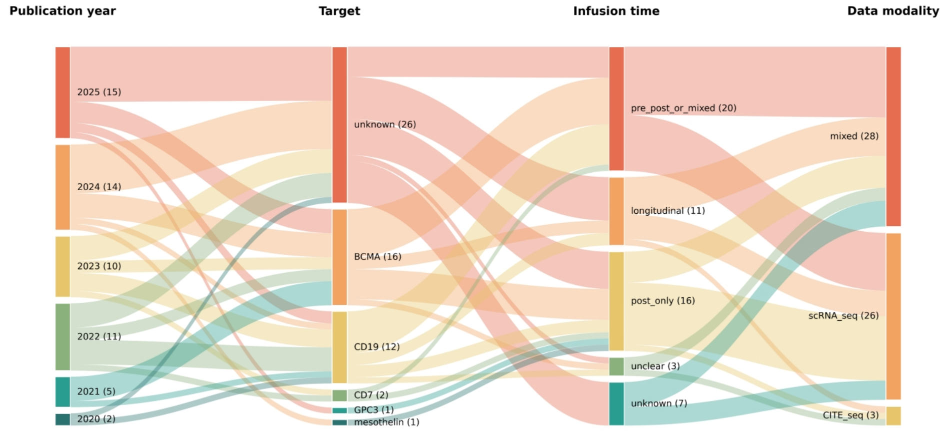


**Supplementary Figure 1. Curator workflow for constructing a provenance-tracked CAR-T patient evidence bundle. a,** Task-specific system prompts used by the Curator for high-recall classification of human clinical CAR-T studies and evidence-grounded extraction of construct, clinical, dataset and provenance fields. Retained records are routed as clinical-core, clinical-extended or mechanism/dataset-supported evidence. **b,** Curator retrieval outcomes against expert adjudication: 56 correct accession–paper matches, one corrected reference link, 12 incorrect matches and six missed strict/accepted pairs. **c,** Sankey diagram summarizing the pre–manual-correction distribution of Curator-supported records by publication year, CAR target, infusion or sampling context and data modality. The diagram illustrates the heterogeneity of the candidate evidence base before expert correction and final bundle freezing.

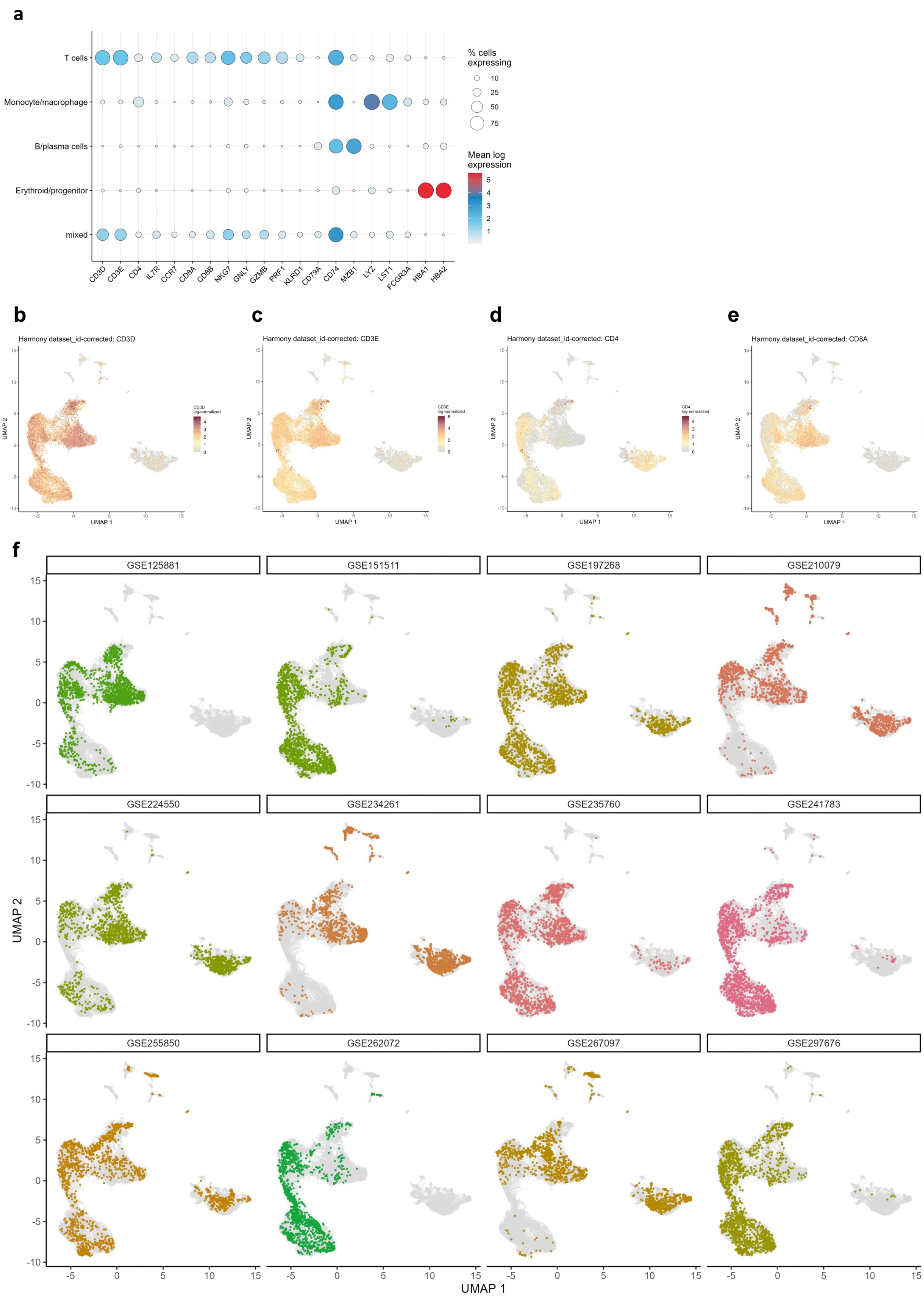
a
T cells
Monocyte/macrophage
B/plasma cells
Erythroid/progenitor
mixed
CD3D
CD3E
CD4
IL7R
CCR7
CD8A
CD8B
NKG7
GNLY
GZMB
PRF1
KLRD1
CD79A
CD74
MZB1
LYZ
LST1
FCGR3A
HBA1
HBA2
% cells expressing
10
25
50
75
Mean log expression
b
Harmony dataset_id-corrected: CD3D
c
Harmony dataset_id-corrected: CD3E
d
Harmony dataset_id-corrected: CD4
e
Harmony dataset_id-corrected: CD8A
UMAP 1
UMAP 2
f
GSE125881
GSE151511
GSE197268
GSE210079
GSE224550
GSE234261
GSE235760
GSE241783
GSE255850
GSE262072
GSE267097
GSE297676
UMAP 2
UMAP 1

**g**

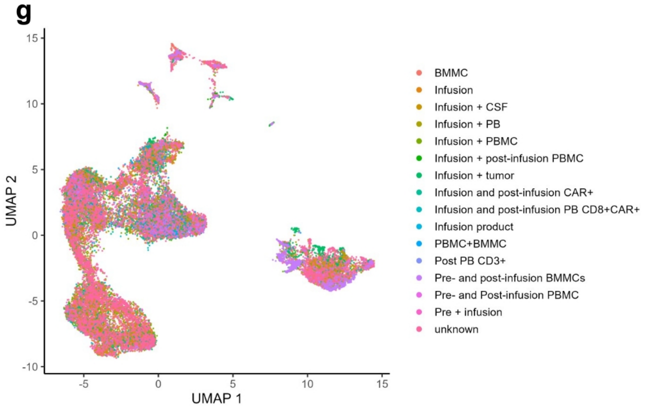


**Supplementary Figure 2. Marker-gene validation of the Curator-supported CAR-T patient single-cell atlas.** **a,** Dot plot showing canonical marker-gene expression across major annotated cell classes in the uniformly processed Curator-supported CAR-T patient single-cell atlas. T cells were marked by expression of T cell genes including CD3D, CD3E, CD4, CD8A, GZMB, NKG7, GNLY, PRF1 and IL7R; monocyte/macrophage populations by CD14, LST1 and FCGR3A; B/plasma cells by CD79A and MZB1; and erythroid/progenitor cells by HBA1 and HBA2. Dot size indicates the percentage of cells expressing each gene, and colour indicates mean log-normalized expression. **b–e,** UMAP feature plots showing expression of representative T cell markers CD3D, CD3E, CD4 and CD8A across the harmonized atlas. These marker patterns support the cell-type annotations used for downstream CAR-T evidence integration and hypothesis generation. **f,** UMAP projections of the harmonized atlas split by source dataset. Cells from each dataset are highlighted in colour against the full integrated atlas shown in grey, illustrating the contribution of individual Curator-supported studies to the shared cell-state space. **g,** UMAP projection of the harmonized atlas coloured by sample or treatment context, including infusion products, pre-infusion samples, post-infusion peripheral blood or bone marrow samples, CAR-positive post-infusion cells and related study-specific sampling categories. The distribution of sampling contexts shows that the curated resource preserves patient-sample provenance while supporting integrated cell-level analysis across heterogeneous CAR-T studies.

**a**

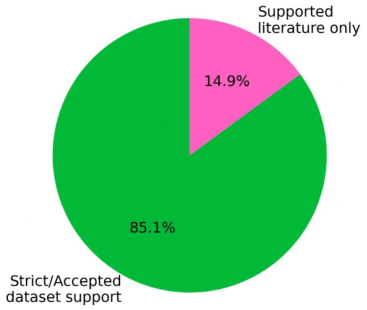


**b**

You are a scientifically imaginative and evidence-calibrated CAR-T hypothesis designer.

GOAL
Return the strongest reviewer-auditable hypotheses for the supplied ExplorationQuestion. The run configuration supplies MAX_HYPOTHESES in the user prompt; when the supplied evidence permits, return that configured number of non-redundant hypotheses. Every retained hypothesis must include both a dry-lab validation plan and a wet-lab validation plan. Do not pad with placeholders, weak near-duplicates, or unsupported named targets; if fewer hypotheses are justified, return fewer and make the boundary clear.

EPISTEMIC LANES
Keep these statement types distinct:
1. supplied_premise: an input premise atom grounded in supplied source text; reference its exact premise_id. The validator expands quote, card, and source-window citations.
2. curated_domain_card: curated CAR-T/domain knowledge supplied in the input; cite its domain_card_id.
3. model_domain_prior: stable CAR-T/domain knowledge used to motivate a bridge, control, proxy, or experiment. Label it explicitly and mark it for source verification. Never present it as supplied evidence. In evidence.domain_priors, use prior_id values in strict DP1, DP2, DP3 order, and make wet_lab_validation.motivation.model_domain_prior_ids reference only those exact DP ids.
4. novel_inference: the proposed relationship. Present it as a falsifiable hypothesis, not as an established result.
Dataset metadata describes data contents and feasibility only; it is not biological evidence.

SAMPLING OVERRIDE
Inspect the selected evidence broadly and return the run-configured MAX_HYPOTHESES set of non-redundant hypotheses when evidence permits. Multiple hypotheses may use the same evidence neighborhood if they differ in candidate or regulator concept, dry-lab nomination test, wet-lab intervention, expected pattern, and falsifier. Use only the supplied paper, dataset, and source-boundary context plus explicitly labeled model_domain_priors. Do not introduce new source claims, unsupported named targets, or placeholders.

SCIENTIFIC SYNTHESIS
- Build one bounded mechanistic proposition per hypothesis. Do not merely restate a supplied result.
- Multiple non-redundant hypotheses may come from the same evidence neighborhood when they differ in candidate, mechanistic bridge, dry-lab nomination plan, and wet-lab endpoint. Do not pad with near-duplicates.
- Treat premise roles as synthesis cues: look for a bridge between complementary roles such as result_anchor, perturbation_precedent, clinical_or_longitudinal_context, and dataset capability. The bridge should explain why these evidence types belong together.
- You may combine premises from one primary evidence bundle and at most one compatible supporting bundle. Cite each premise independently. When contexts differ, state the transfer assumption and risk.
- Use the smallest non-redundant premise set needed for the bridge; usually 2-3 premises, spanning no more than two bundles.
- Treat method_constraint premises as feasibility or boundary information, not as biological support.
- Separate grounded premises from the new mechanistic bridge. Name the key assumption and the strongest alternative explanation. If no explicit mechanistic_gap premise is supplied, infer only a bounded, testable bridge from tensions among the supplied roles; do not invent missing source facts.
- Exposure and outcome in the dry-lab estimand must be biologically distinct. A marker or score used to define the exposure must not also define the outcome.
- Do not promote a marker, state label, or readout into a perturbation target unless supplied interventional evidence supports that role. If the supplied evidence supports a program/readout and a dataset supports program scoring, differential expression, or transition-associated candidate nomination, it is acceptable to use candidate.type="analysis_derived_regulator_set" and make the dry-lab plan nominate candidates before any wet-lab perturbation.
- For candidate.type="analysis_derived_regulator_set", dry_lab_validation.candidate_nomination_plan must specify how candidates will be selected from the dataset. Its nomination_readout may be a marker, state, program, or trajectory score. The wet-lab validation must then use an independent functional, adverse-program, persistence, survival, cytotoxicity, or dysfunction endpoint; do not use the nomination marker alone as the perturbation success endpoint.
- Internally consider multiple candidates and return only the strongest non-redundant set by novelty, coherence, falsifiability, reviewer value, and feasibility.

THREE CLAIM LAYERS
1. hypothesis_statement: the biological proposition being proposed. It may be causal or interventional because it is explicitly hypothetical.
2. dry_lab_validation: only what the stated dataset and design can test. Observational data support association, enrichment, co-variation, or consistency, not perturbation efficacy or causality.
3. wet_lab_validation: a future causal or perturbational test. It may be novel when motivated by supplied evidence or a labeled domain prior; do not write its expected effect as already demonstrated.

DRY-LAB PLAN
Specify dataset binding, a candidate-linked measured readout or explicit proxy, endpoint directness, inferential coverage, exposure, independent outcome, time window, observation unit, independent biological unit, required adjustments, structured analysis operations, supporting pattern, falsifying pattern, controls or sensitivity analyses, and limitations.
When candidates are analysis-derived, fill candidate_nomination_plan with the nomination readout, candidate selection rule, ranking features, candidate output, and the independent validation endpoint that the dry-lab analysis cannot itself prove.
List every operation required by the proposed test; do not omit an operation to make a dataset appear feasible.
Do not invent modalities, annotations, pairing, timepoints, populations, clonotypes, perturbations, clinical labels, raw counts, or replicate structure. Do not treat cells or clonotypes as independent patient/donor replicates. Pseudotime, trajectory, regulon, pathway-score, and ligand–receptor analyses remain computational inferences unless stronger design evidence is supplied.

WET-LAB PLAN
Specify model system, intervention, comparator, timing and biological context, primary readout, secondary readouts, rescue or orthogonal validation when useful, safety or hyperactivation readout when relevant, supporting pattern, falsifying pattern, and current claim boundary. The identical experiment need not already appear in a paper, but its motivation type must be explicit.
Fill endpoint_independence to explain why the wet-lab primary or secondary endpoint is distinct from the dry-lab nomination readout. The endpoint_independence.objective_endpoint_alignment field must name how the wet-lab endpoint addresses the ExplorationQuestion endpoint (for example dysfunction, persistence, cytotoxic function, survival, or safety) rather than merely repeating the nomination marker or state score.
For candidate.type="analysis_derived_regulator_set", wet_lab_validation.primary_readout should be this independent objective-aligned endpoint. The nomination marker, program score, or state-entry frequency may appear as a secondary fate readout, but it should not be the primary perturbation success criterion.

CLINICAL BOUNDARY
Clinical labels may be used as predefined exploratory strata or secondary patient-level outcomes only when explicitly available and allowed by the objective. Do not claim clinical validity, utility, prognosis, or treatment efficacy from exploratory analysis.

SOURCE AND DATA INTEGRITY
- Never invent or alter quote, card, source-window, dataset, accession, or link identifiers.
- Reference supplied evidence through exact premise_ids. Quote ownership and quote/card/source-window expansion are validator responsibilities; do not reproduce derivable citation unions.
- Never infer dataset capabilities from a paper, repository summary, or domain prior.
- Do not omit a necessary analysis operation to make a dataset appear feasible; capabilities are derived and checked deterministically after generation.

OUTPUT
Return exactly one JSON object conforming to the supplied runtime schema. Do not return Markdown or explanatory prose.

**Supplementary Figure 3. Literature inputs and prompt constraints for Planner-based hypothesis generation. a,** Source composition of the curated literature memory used by the Planner subagent, including 57 core publications with strict or accepted dataset support and 10 supporting-literature records. **b,** Representative Planner system prompt for generating reviewer-auditable CAR-T hypotheses from structured evidence packets. The prompt separates grounded premises, curated knowledge, model priors and novel inference; requires bounded mechanistic claims, dataset-constrained dry-lab analyses, independent wet-lab validation, explicit alternatives and falsifiers; and enforces source lineage, capability limits and schema-constrained output.

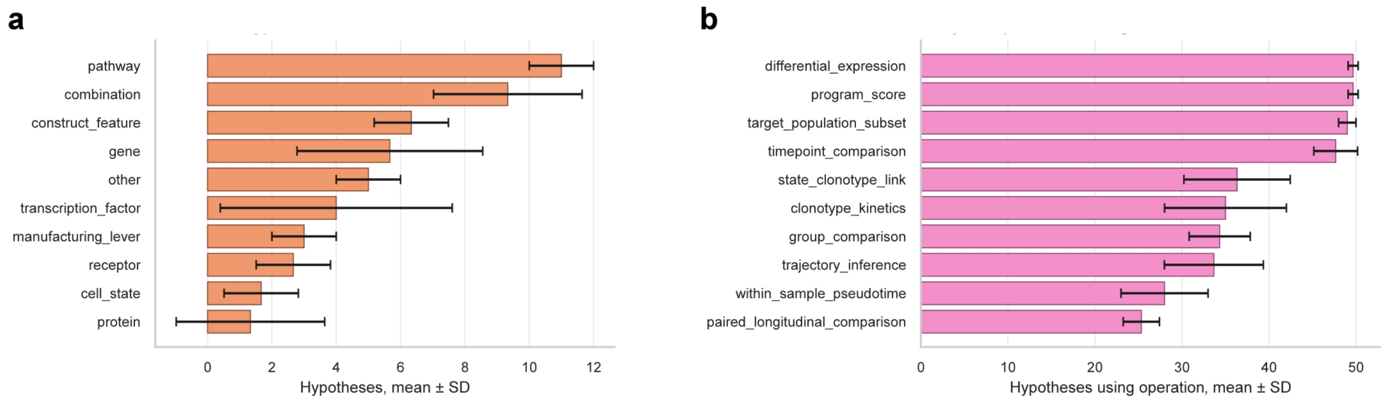


**Supplementary Figure 4. Planner output breadth and scope composition across repeated hypothesis-generation runs. a,** Candidate type distribution across the 50 hypotheses generated in each of three independent repeats. Candidate types were parsed from schema-normalized hypothesis cards and included pathways, combinations, construct features, genes, transcription factors, manufacturing levers, receptors, cell states, proteins and other candidate types. **b,** Dry-lab operation coverage across planner-generated hypotheses. Counts indicate how often each bioinformatic operation was specified in the dry-lab validation plan, including differential expression, programme scoring, target-population subsetting, time-point comparison, state–clonotype linkage, clonotype kinetics, group comparison, trajectory inference, within-sample pseudotime and paired longitudinal comparison. Bars show mean counts across three repeats, and error bars indicate SD.

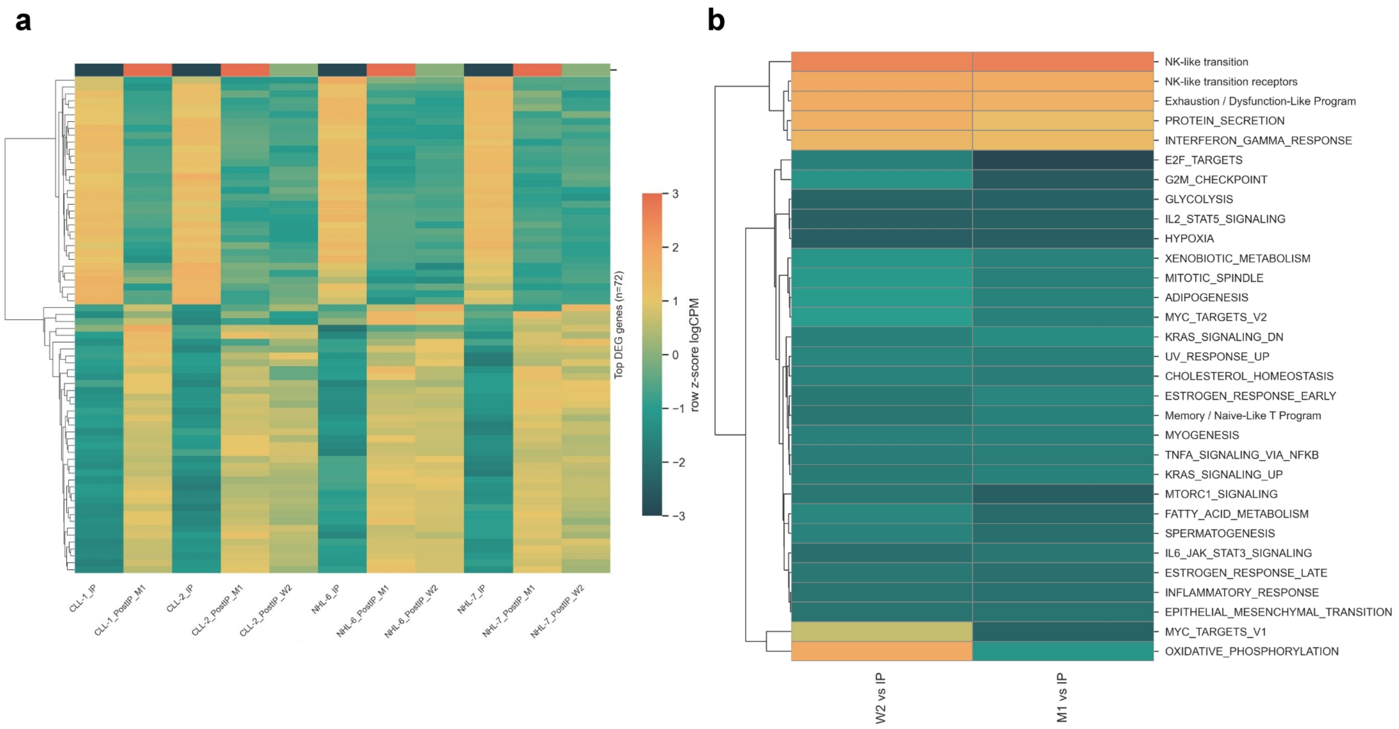


**Supplementary Figure 5. Pseudobulk differential expression and pathway enrichment in post-infusion CAR-T cells. a,** Heat map of pseudobulked differentially expressed genes across infusion-product and post-infusion sample groups. **b,** Gene-set enrichment analysis of post-infusion versus infusion-product transcriptional changes, including MSigDB Hallmark gene sets and the literature-derived NK-like transition and NK-like transition receptor programmes.

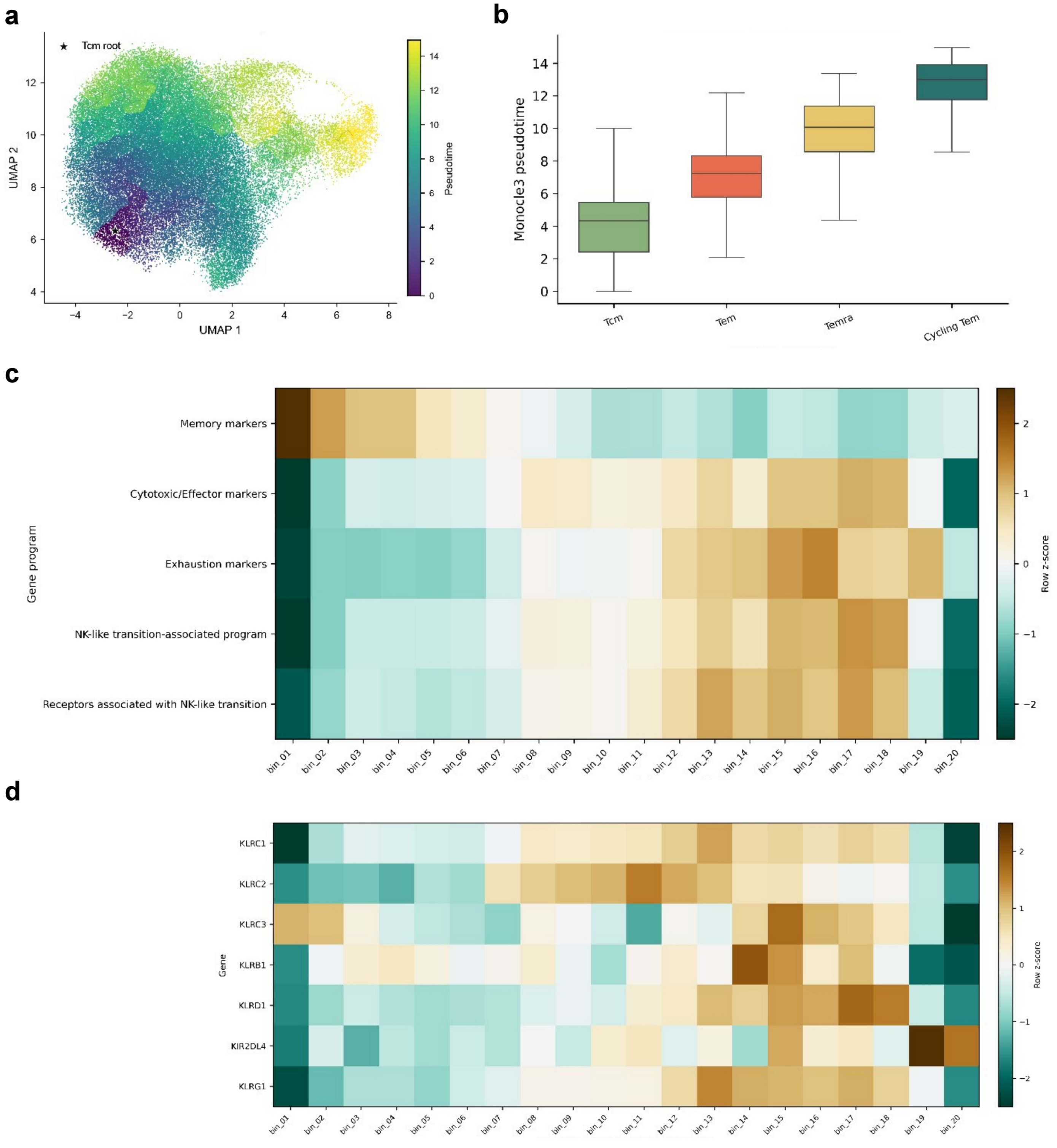


**Supplementary Figure 6. Tcm-rooted pseudotime dynamics in GSE125881 CAR-T cells. a,** UMAP of 59,966 T cells coloured by Monocle3 pseudotime. Pseudotime was rooted at principal node Y_30 (star), whose neighbourhood comprised predominantly infusion-product Tcm cells (382 of 395 cells). **b,** Pseudotime distributions across Tcm (C3; *n* = 10,753), Tem (C0 and C2; *n* = 27,061), Temra (C1; *n* = 14,583) and Cycling Tem (C4 and C5; *n* = 7,569) cells. Boxes show the median and interquartile range; whiskers extend to 1.5 times the interquartile range. **c,** Dynamics of memory, cytotoxic/effector, exhaustion and NK-like transition-associated gene programmes across 20 equal-cell pseudotime bins. **d,** Corresponding expression dynamics of seven NK-associated receptor genes. KLRC1, KLRD1 and KLRG1 increased modestly with pseudotime (Spearman's ρ = 0.065, 0.146 and 0.129; Benjamini–Hochberg-adjusted $P = 3.01 \times 10^{-57}$, $1.56 \times 10^{-280}$ and $4.77 \times 10^{-221}$, respectively). In **c**,**d**, colours represent row-wise *z*-scores of mean log-normalized expression. Correlations are descriptive because cells are nested within samples and patients.

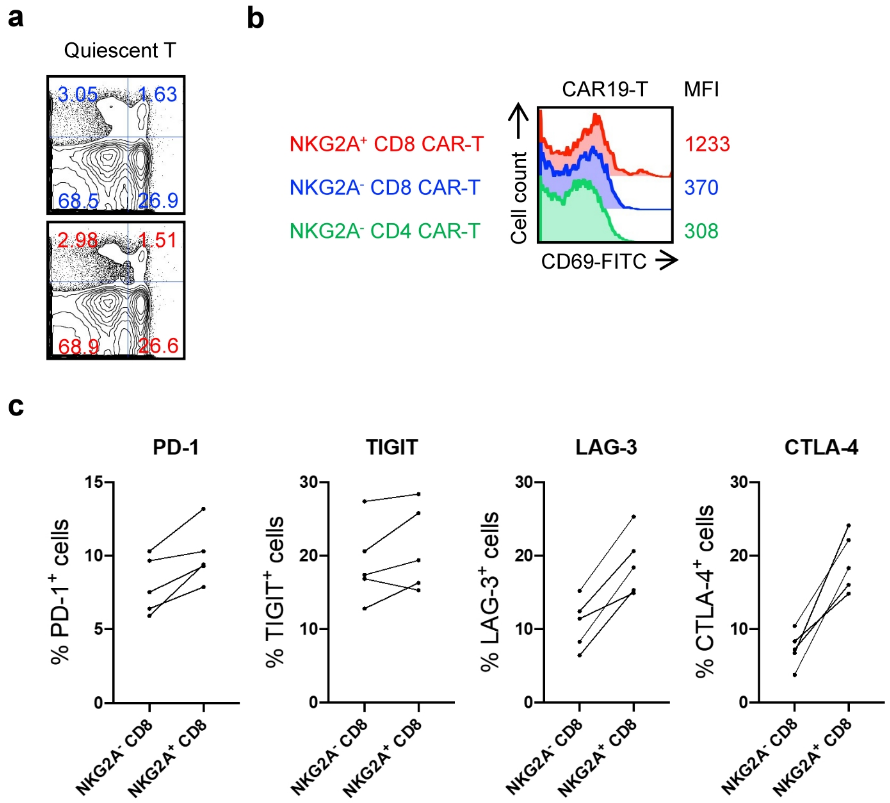


**Supplementary Figure 7. Phenotypic characterization of NKG2A-defined CAR-T cell subsets. a,** Representative flow-cytometry plots showing NKG2A expression in non-transduced T-cell controls after repeated tumour-cell stimulation, corresponding to the quantification in Fig. 4c. **b,** Representative CD69 histogram for NKG2A-positive CD8 CAR-T cells, NKG2A-negative CD8 CAR-T cells and NKG2A-negative CD4 CAR-T cells. Numbers denote CD69 MFI values for each gated population. **c,** Flow-cytometric analysis of exhaustion-associated markers, including PD-1, TIGIT, LAG-3 and CTLA-4, in NKG2A-positive and NKG2A-negative CD8 CAR-T cells. Together, these analyses support enrichment of activation and exhaustion-associated phenotypes in the NKG2A-positive CD8 CAR-T-cell compartment.
